# Multimodal imaging reveals a lysosomal drug reservoir that drives heterogeneous distribution of PARP inhibitors

**DOI:** 10.1101/2025.05.31.656628

**Authors:** Carmen R. Moncayo, Restuadi Restuadi, Guanying Zhang, Daniel Marks, Paula Ortega-Prieto, Emily Doherty, Nathalie Lambie, Chad Whilding, Ivan Andrew, Alex Montoya, Bhavik Patel, Betheney R. Pennycook, Vincen Wu, Zoltan Takats, Nik Matthews, George R. Young, Pavel Shliaha, Laurence Game, Boris Lenhard, Iain McNeish, Christina Fotopoulou, Alexis R. Barr, Paula Cunnea, Zoe Hall, Louise Fets

## Abstract

For all drugs, effective target engagement requires sufficient intracellular concentrations of drug to be reached, but whether tumour heterogeneity impacts drug distribution and efficacy is poorly studied. PARP inhibitors have transformed treatment of high-grade serous ovarian carcinoma (HGSOC), but resistance remains a clinical hurdle in this highly heterogeneous tumour type. We developed a patient-derived explant multi-modal imaging pipeline, which demonstrated that cell-intrinsic PARP inhibitor accumulation is highly variable, both between patients and within tumours. Spatial transcriptomics revealed enrichment of apoptotic and lysosomal signatures in ‘high-drug’ regions. Rucaparib, an intrinsically fluorescent PARP inhibitor, accumulates heterogeneously at the single-cell level, with ‘rucaparib-high’ cells demonstrating increased drug response relative to ‘rucaparib low’. Mechanistically, lysosomal sequestration creates a rucaparib reservoir that determines drug levels in the nucleus. Perturbation of lysosomal content altered intracellular levels of weak base PARP inhibitors rucaparib and niraparib, but not olaparib. Together these data suggest that lysosomes act as a reservoir for a subset of PARP inhibitor drugs to improve drug response.

## Introduction

To be effective in tumours, therapeutic drugs must reach a concentration at which they can effectively bind their target, and the intracellular concentration achieved will also dictate off-target effects^15^. Though it is clear that tumour heterogeneity can drive therapeutic resistance^1^, both at the inter-patient level and at the intra-tumoural level, whether tumour heterogeneity impacts upon drug accumulation and distribution, and how this influences drug efficacy and resistance has yet to be explored.

High grade serous ovarian carcinoma (HGSOC) is a particularly heterogeneous disease at the intra-patient level, both spatially and temporally^2,3^. Genetic heterogeneity is well described, and includes near-ubiquitous *TP53* mutations^4^, substantial copy number variation and high-frequencies of homologous recombination DNA repair deficiency (HRD), most commonly through *BRCA1/2* mutation^5–7^. Non-genetic heterogeneity is also well documented in HGSOC, even within fallopian tube epithelia, the cell type of origin^8^, and substantial phenotypic diversity has been observed in tumour cells from multiple patient cohorts^3,9–11^.

Patients with HGSOC typically undergo cytoreductive surgery, combined with platinum-based chemotherapy. The introduction of additional targeted maintenance therapies such as PARP inhibitors has led to significantly improved remission rates in HGSOC patients^12,13^. PARP inhibitors exploit HRD and lead to synthetic lethality, with three agents in current clinical use for HGSOC: olaparib, niraparib and rucaparib. Mechanistically, these drugs cause PARP molecules to become trapped on DNA in addition to inhibiting their catalytical activity. This results in replication fork stalling and the formation of DNA double strand breaks (DSBs)^14^, which ultimately lead to apoptosis of tumour cells.

Variability in PARP inhibitor accumulation, at the intra- or inter-patient level, has yet to be studied, despite evidence that fluorescent analogues of PARP inhibitors accumulate heterogeneously in *in vitro* and pre-clinical *in vivo* models^16^. Whether this is mimicked by the clinically used drugs is unknown, but could be of significant consequences if a sub-population of cells accumulates drug to a level below that required for target engagement, which could contribute to resistance. Therefore, we set out to determine whether PARP inhibitors are heterogeneously accumulated in HGSOC and whether this impacts their efficacy.

Here, using HGSOC patient-derived explant (PDE) models, we developed a pipeline to treat tumour slices with PARP inhibitors *ex vivo*. Mass spectrometry imaging (MSI) demonstrated spatial heterogeneity in drug distribution both between and within patient tumours. We used spatial transcriptomics of adjacent tissue sections to compare regions of ‘high-’ and ‘low-drug’ concentration, revealing that differential drug accumulation has significant effects on drug response and is associated with lysosomal signatures. Using established ovarian cancer cell lines, we demonstrate that heterogeneity in drug response occurs at the single-cell level, and that this is linked to cell-to-cell variability in drug accumulation. Mechanistically, we demonstrate that lysosomal sequestration of both rucaparib and niraparib, but not olaparib, drives increased uptake in ‘high-drug’ cells. Together, these data suggest that heterogeneous drug distribution may be a contributing factor to drug resistance in a clinical setting.

## Results

### A vasculature-independent, spatially-resolved pipeline to assess PARP inhibitor distribution in Patient-Derived Explants

We hypothesised that the substantial genetic and non-genetic heterogeneity in HGSOC^2,9,10^ may impact distribution of PARP inhibitors within tumours. To test whether cell-intrinsic properties could lead to differential drug accumulation, we required a model in which PARP inhibitors could be delivered to the tumour independently of the vasculature. To achieve this, we used a PDE model using surplus surgical tissue from treatment-naïve patients with HGSOC who had undergone maximal-effort primary cyto-reductive surgery. Samples from five tumours, derived from three patients, were each sectioned into multiple slices (details in Supplementary Data Table 1) and cultured *ex vivo*, enabling us to incubate slices from the same tumour with three PARP inhibitors; niraparib, olaparib and rucaparib. Twenty one independent PDEs were processed using this pipeline in total. After treatment, each PDE slice was snap frozen, then cryo-sectioned into multiple 10 µm-thick sections to assess drug distribution, drug response and tissue morphology (Fig. 1a).

**Figure 1:**
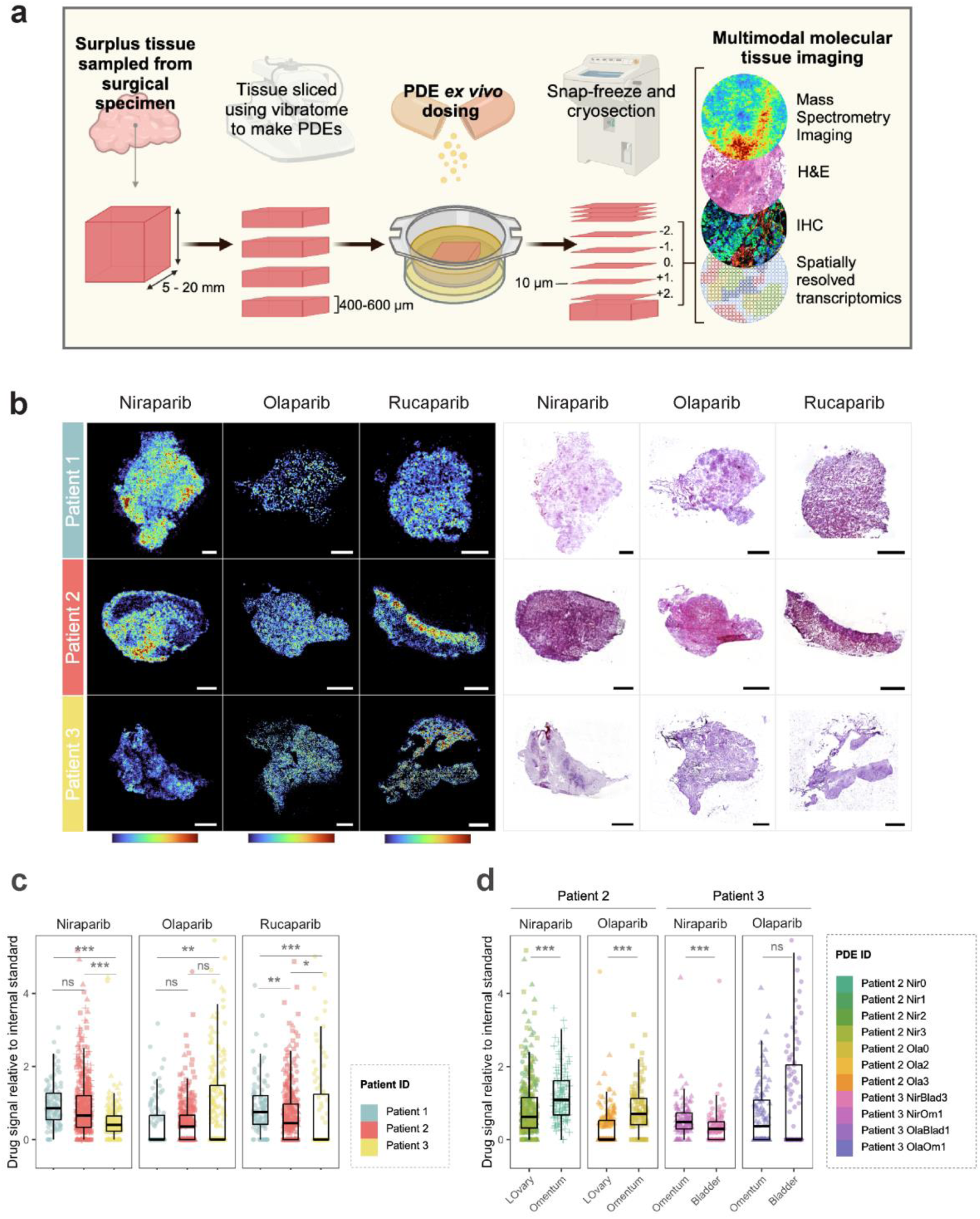
Mass Spectrometry Imaging reveals vasculature-independent inter- and intra-patient heterogeneity in PARP inhibitor distribution. A) A vasculature independent, patient-derived explant (PDE) pipeline to investigate mechanisms of drug accumulation, response and resistance in a spatially-resolved manner. H&E: haematoxylin and eosin; IHC: immunohistochemistry. B) Representative MALDI mass spectrometry images (MSI) of PARP inhibitor drug distribution in PDEs dosed ex vivo. Niraparib [M+Na]^+^ (343.1535 ± 5 ppm), olaparib [M+Na]^+^ (457.1652 ± 5 ppm) and rucaparib [M+H]^+^ (324.1512 ± 10 ppm) signals, relative to drug-matched internal standards, are shown on the left. To the right, H&E stainings of the same tissue sections post-MSI. Scale bar =1 mm C) Inter-patient heterogeneity in PARP inhibitor accumulation. Drug signal (per pixel, relative to matched internal standards) was normalised to the third quartile (Q3) of combined data from all samples within that drug, to enable plotting of all three drugs on the same scale. Tissue pixels were grouped into percentiles based on drug distribution, with each percentile represented as an individual data point. Technical replicate PDEs are plotted with different shapes. Wilcoxon rank sum test with Bonferroni correction was applied over percentiles to compare differences in drug signal intensities across patients. D) Inter-tumour heterogeneity in PARP inhibitor accumulation. Data was processed as described in (C).

To visualise drug distribution in the treated-PDE tissue sections, we used mass spectrometry imaging (MSI), which showed excellent signal linearity with concentration for all three drugs tested (Extended Data Fig. 1a-c). Using olaparib-treated samples as a test-case, MSI was used to image drug distribution in slices treated for varying amounts of time, from which we determined that steady state drug distribution was reached after 24h of treatment (Extended Data Fig. 1d-f).

The distributions of niraparib, rucaparib and olaparib were compared in 18 PDEs cultured after tumour resection from three patients with HGSOC. One additional PDE from each patient was treated as a vehicle-only control to check the specificity of the MSI signal. PARP inhibitor signal was normalised relative to a heavy isotope-labelled analogue of each drug, homogeneously deposited across each tissue slice as an internal standard, to enable semi-quantitative comparison of drug accumulation between patients. Where tissue availability allowed, multiple slices were cultured for each patient/drug combination, and tumours were sourced from multiple sites (Supplementary Data Table 1). For all tissue slices, si× 10 µm-thick sections were used for multi-modal imaging analyses. The section analysed using MALDI MSI, deemed ‘section 0’, was subsequently haematoxylin and eosin (H&E) stained to examine tissue architecture. The adjacent section, ‘section +1’ was used for GeoMx spatial transcriptomics, while immunohistochemical (IHC) staining of section −2, −1, +2 and +3 probed apoptosis and DNA damage levels using cleaved caspase 3 and ɣH2AX, as well as HGSOC markers Wilms’ tumour 1 (WT1) and Pax8. Extended Data Figure 2 presents the results of these analyses in a grid form; specific comparisons will be referred to using the coordinates of this grid. FIGO stages of patients can be found in Supplementary Data Table 1.

### Cell-intrinsic heterogeneity in PARP inhibitor accumulation in HGSOC explants

Despite dosing independently of the vasculature system, a high degree of heterogeneity in PARP inhibitor distribution was observed across all patients (Fig. 1b), with clear drug ‘hotspots’ in some parts of the tumour, while other regions accumulated little or no detectable drug. Interestingly, the uptake of the three PARP inhibitors varied between patients, with relative levels of niraparib and rucaparib being comparable in the different patients, while olaparib accumulation differed (Fig. 1b, c). For example, samples from Patient 1 accumulated very little olaparib, whereas levels of niraparib and rucaparib were highest in these samples relative to the others tested. This suggests that the differential uptake of drugs between patients is not due to a general difference in small molecule permeability within these tumours, but that specific mechanisms may govern the uptake and/or efflux of rucaparib and niraparib when compared to olaparib.

Although some examples of this heterogeneity were associated with differences in tissue architecture by H&E staining (e.g., a correlation between drug intensity in regions of increased cell density in Patient 2 after treatment with niraparib or rucaparib, Extended Data Fig. 2, E3:4,N3:4), this was not always the case (e.g. Patient 1, niraparib, A3:4). Regions of ‘high’ and ‘low’ drug accumulation did not appear to correlate with levels of tumour markers such as Pax8 or WT1 (Extended Data Fig. 2, rows 3, 9 and 10), suggesting that the drug does not exclusively accumulate in cancer cells, and within the cancer cell populations, there is variability in accumulation. In a number of cases, there was a correlation between the drug distribution and the staining pattern of ɣH2AX, a readout of DNA damage levels (Extended Data Fig. 2b, rows 2 and 3).

Inter-tumour heterogeneity in drug accumulation was also observed between different tumours from the same patient (Fig. 1d, Extended Data Fig. 2 row 3). Niraparib-treated samples from patient 2 derived from the omentum (E) and left ovary (F,G,H) showed differing drug accumulation, as did olaparib-treated samples derived from the same sites (I and J,K respectively). Similarly, niraparib-treated samples derived from the bladder (P) and omentum (Q) of patient 3 displayed variability, as did olaparib-treated samples from these locations (R, S). Together, these data demonstrate that the accumulation of PARP inhibitors in patient-derived explants is highly heterogeneous, at the intra-tumour, inter-tumour and inter-patient level.

### Regions of low drug uptake show significantly lower drug response signatures than drug hotspots

To understand what drives the differential accumulation of PARP inhibitors and how this impacts their efficacy, we used the Nanostring GeoMx spatial transcriptomics platform to determine whole-transcriptome differences between ‘high-’ and ‘low-drug’ regions of our PDEs. To achieve sufficient sequencing depth for whole-transcriptome analysis using this platform, a minimum number of nuclei are recommended per ROI. This was difficult to achieve in olaparib-treated samples as drug hotspots were, on average, much smaller than those observed in niraparib or rucaparib-treated samples (Fig. 1b, Extended Data Fig. 2, row 3), therefore we chose to focus only on the latter two drugs.

Regions of ‘high-’ and ‘low-drug’ were chosen based on MSI images of drug distribution and cross-referenced with the tissue morphology marker pan-cytokeratin and α-smooth muscle actin, to confirm that transcriptional profiles obtained were from cancer cells only and that fibroblasts were excluded (Fig. 2a, Extended Data Fig. 2, rows 5:8). Finally, regions were compared with IHC staining of ovarian cancer cell-specific markers, WT1 and Pax8, on alternative sections (Extended Data Fig. 2, rows 9:10). Twelve tissue sections were analysed, and an average of 5 ‘high-’ and 5 ‘low-drug’ regions of interest (ROIs) were obtained per tissue section. Quantification of the levels of drug between the ROIs confirmed a significant increase in drug levels in ‘high’ ROIs relative to ‘low’ for both niraparib and rucaparib (Fig. 2b).

**Figure 2:**
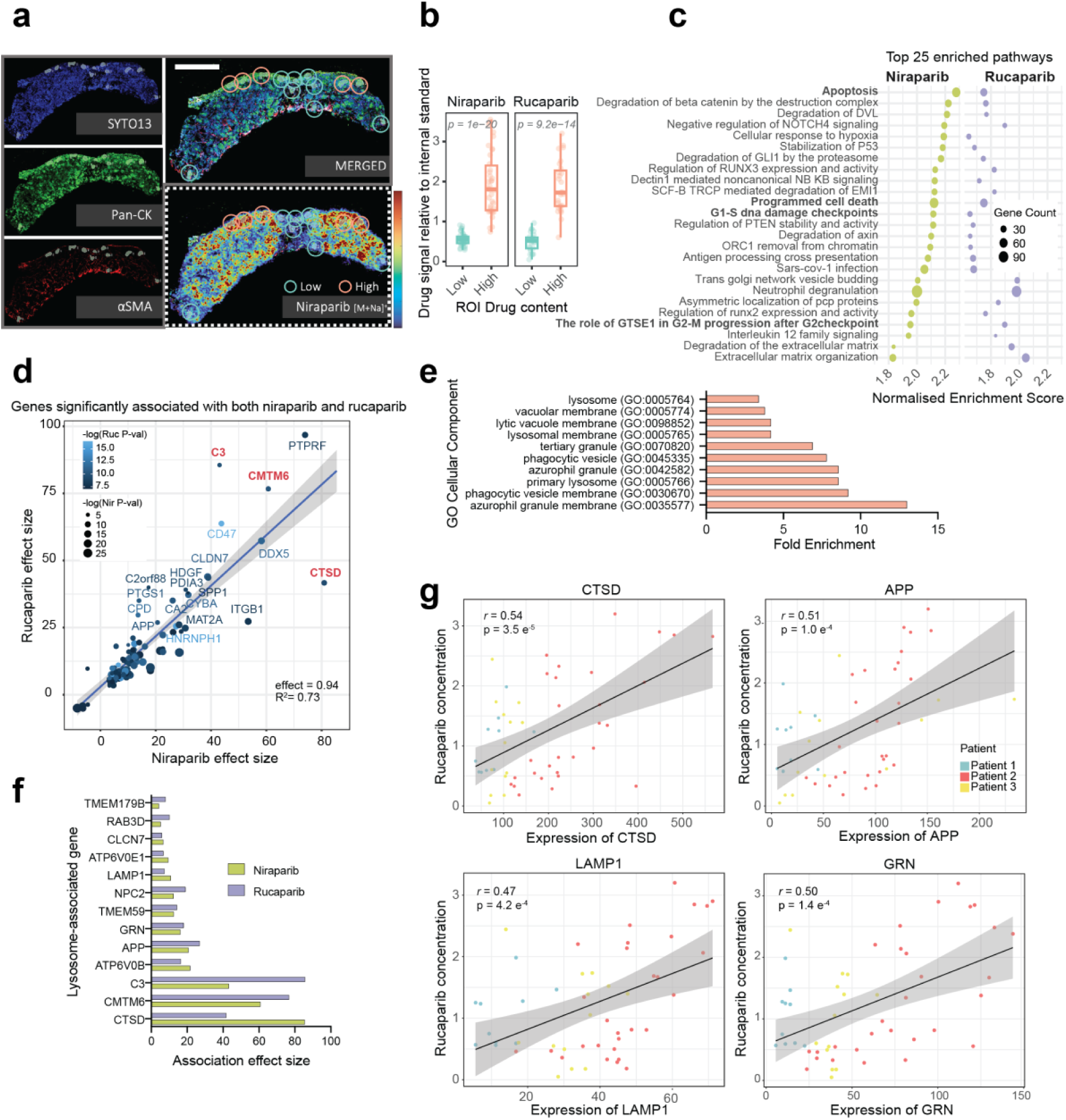
Regions of high PARP inhibitor accumulation show increased apoptotic response and lysosomal signatures. A) Example of ‘high’ and ‘low’ region of interest (ROI) selection for GeoMx spatial transcriptomics in PDEs following ex vivo dosing with PARP inhibitors. The MSI image in the dashed bottom-right panel, displaying niraparib distribution, was used to inform ROI selection on panCK-positive, αSMA-negative tumour areas. Scale bar = 1mm. B) ROI drug signal intensity quantification, relative to internal standards spiked in matrix, and Q3-normalised (as in 1C) to enable same-scale plotting. Wilcoxon rank sum test was applied to evaluate differences in drug intensity signal between ‘low’ and ‘high’ ROIs. C) Top 25 Reactome pathways enriched in both niraparib (green) and rucaparib (purple) datasets following Gene Set Enrichment Analysis (GSEA). D) Correlation of effect sizes for genes with expression significantly associated with both niraparib and rucaparib levels per ROI. Drug levels were modelled as continuous variables, derived from mean signal intensities per ROI in mass spectrometry images, normalised to heavy-labelled drug-analogue internal standards. Associations were estimated using a linear mixed model accounting for hierarchical structure (patient and PDE as nested random effects). Point size and colour indicate the significance of niraparib and rucaparib effects, respectively. E) Lysosome-related terms identified in a GO Cellular Compartment analysis of genes commonly associated with increasing levels of niraparib and rucaparib. F) Effect sizes of lysosomal genes significantly associated with increasing niraparib (green) and rucaparib (purple) levels in PDE samples. G) Scatter plots depicting expression of lysosomal genes relative to rucaparib concentration from PDE experiments, each dot represents an ROI and patients are colour-coded. Pearson correlation coefficients indicated.

After QC and filtering steps, we were left with 122 ROIs, with at least 20 regions of ‘high-’ and ‘low-drug’ for each of the two PARP inhibitors (Extended Data Fig. 3a), and across these, 11224 targets were reliably quantifiable after filtering. PCA and Euclidean distance clustering analysis demonstrated that, as expected, the major driver of transcriptional variance was inter-patient heterogeneity (Extended Data Fig. 3b, c). Within each patient, samples grouped by compound, then by PDE slice ID, and within each slice, ‘high’ and ‘low’ drug content areas were separated from one another, demonstrating that differential drug accumulation was associated with divergent transcriptomes (Extended Data Fig. 3b). Pathway analysis of all ‘high’ versus ‘low’ regions using the Reactome database demonstrated that ‘high-drug’ regions were enriched in terms associated with cell cycle arrest and DNA damage response (‘G1/S Damage Checkpoint’), and apoptosis (‘Apoptosis’, ‘Apoptotic execution phase’, ‘Programmed cell death’, ‘Apoptotic cleavage of cellular proteins’), suggesting that PARP inhibitor response was enhanced by the increased concentration of the drugs, driving increased apoptotic signalling (Fig. 2c, Supplementary Data Table 2).

### High drug accumulation is associated with lysosomal gene signatures

By incorporating the relative drug concentrations quantified across each ROI (Fig. 2b), we were able to use drug level as a continuous (rather than a binary ‘high’ versus ‘low’) variable as part of a separate linear mixed model for each of our two drugs used, with nested random effects to account for patient and PDE slice-derived variability. Using this, we identified genes whose expression was significantly associated with increasing, or decreasing levels of either rucaparib or niraparib (Extended Data Fig. 4a, b, Supplementary Data Table 3). Enrichment analysis identified several overlapping terms associated with increasing levels of both drugs, centring around proteolysis or peptide activity, extracellular matrix features and glycolytic activity (Extended Data Fig. 4c, Supplementary Data Table 4).

Though far more significant gene associations were observed with increasing niraparib concentration (2541, <0.05 FDR), 55% of genes that were associated with increasing rucaparib concentration (201, <0.05 FDR) were also found to be associated to niraparib (111 labelled gene transcripts, Fig. 2d, Data Table 3). Strikingly, the effect sizes of transcripts that were significantly associated with both drugs were highly correlated (Fig. 2d, slope = 0.94, *R*^2^ = 0.73).

Among the transcripts most strongly associated with increasing concentrations of niraparib and rucaparib were *CTSD*, encoding the lysosomal protease cathepsin D; the complement component C3, which has been demonstrated to be found in lysosomes in several cell types^17,18^; and *CMTM6*, the protein product of which has been shown to stabilise PD-L1 at the plasma membrane by promoting its sorting towards recycling endosomes and away from lysosomes^19,20^ (Fig. 2d, relevant genes highlighted in red). Mirroring these associations with the endosome/lysosome system, GO term Cellular Component enrichment analysis of the genes commonly associated with both drugs revealed that ten of the top twenty terms referred to lysosomes or lysosome-related organelles (e.g. azurophilic granules, lytic vacuoles) (Fig. 2e, Supplementary Data Table 5).

On closer inspection 13 of the 111 common gene transcripts encoded lysosome-associated proteins (Fig. 2f), and the correlation between levels of drug and level of transcript expression was striking for well-known lysosomal genes such as LAMP1, CTSD, APP and GRN (Fig. 2g, Extended Data Fig. 4d). We also noted that ‘Neutrophil Degranulation’, a process known to be reliant on the secretion of lysosomal proteases, was also enriched in our binary analysis (Fig. 2c). Together, these data suggest that increased accumulation of both rucaparib and niraparib are associated with lysosomal signatures in PDEs.

### Single cell heterogeneity in PARP inhibitor-induced DNA damage response

PDE models capture HGSOC tumour heterogeneity and cellular diversity well, however 24-hour treatment times are required to reach steady state levels of drug in this system. This prolonged PARP inhibitor exposure prevents the deconvolution of features which drive heterogeneous drug accumulation, from those features that arise as a result of concentration-dependent differences in drug response. We therefore set out to establish a cell-based model to better understand whether high-drug-associated lysosomal signatures could be a cause or a consequence of differential PARP inhibitor accumulation.

To determine whether we could model heterogeneous PARP inhibitor accumulation and response in cell lines, we first examined response to olaparib, rucaparib and niraparib at the single-cell level in established HGSOC cell line, PEO1, using accumulation of ɣH2AX as a read out for DSBs (Fig. 3a). Dosing at their IC_50_ concentration (Extended Data Fig. 5a, b), DNA damage response to PARP inhibitors at the single cell level was highly variable in PEO1 and other ovarian cell lines (Fig. 3b, Extended Data Fig. 5c), as demonstrated previously in other cancer cell types^21^. Separating cells based on DNA content revealed a cell cycle-dependent response, in line with the known mechanism of action of PARP inhibitors. As expected, cells in the G2/M-phase demonstrated higher levels of ɣH2AX compared to vehicle-treated controls. However, even within this subset, a substantial degree of heterogeneity in drug-induced DNA damage was observed.

**Figure 3:**
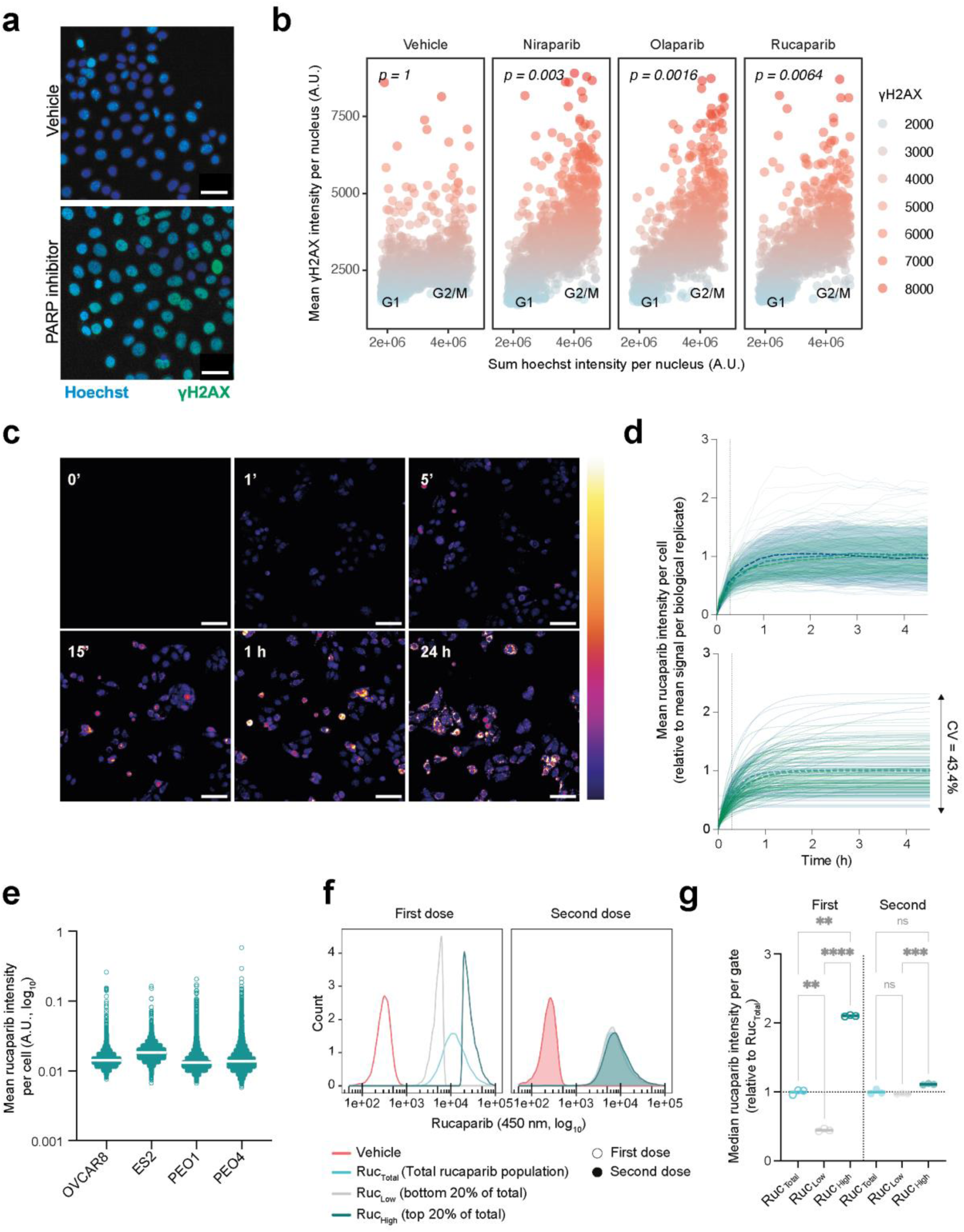
Non-genetic heterogeneity drives variable rucaparib accumulation at the single cell level. A) Representative image of γH2AX levels in PEO1 cells after 24-hour treatment with PARP inhibitors at IC_50_ value. Scale bar = 50 μm. B) Scatter plots, depicting mean γH2AX fluorescence intensities as a function of sum hoechst fluorescence per nucleus upon PARP inhibitor treatment of PEO1 cells. Cell cycle gates were estimated based on hoechst sum fluorescence intensity per nuclei. Data represent n = 3 wells per drug and are representative of 3 biological replicates. Welch two sample t-test over mean γH2AX per technical replicate was applied to compare DNA damage levels upon PARP inhibitor treatment. C) Rucaparib uptake time course in PEO1 cells dosed at IC_50_ concentration. Scale = 25 μm. D) Quantification of rucaparib accumulation across time at the single cell level in PEO1 cells (top). Dotted lines and shaded areas show the mean and standard deviation of rucaparib uptake across 50 cells per biological replicate (n = 3, colour coded). A one-phase association model was fitted (bottom) to estimate uptake rate and steady state rucaparib per cell. E) Heterogeneity in intracellular rucaparib concentration in OvCa cell lines, each dosed at C_max_ of rucaparib (6 µM) for 24h. Population median rucaparib signal is showed as solid horizontal lines. Data is representative of n = 3 biological replicates. F) Phenotypic plasticity of differential rucaparib accumulation. Ruc_High_ and Ruc_Low_ PEO1s (top and bottom 20% of rucaparib-treated population respectively) were FACS-sorted with total drug and vehicle-only controls after a 2-hour treatment. Cells were re-cultured in drug-free medium for 21 days and then re-dosed to analyse rucaparib signal distribution, with respect to its original distribution. Data from n = 3 technical replicates; see Extended Data Fig. 6c for biological replicates of cells re-cultured for 1 and 2 weeks. G) Relative drug levels for each rucaparib population in (F). ‘First’ denotes initial rucaparib signal. ‘Second’ refers to re-dosed levels after re-culture. Data from n = 3 technical replicates; see Extended Data Fig. 6c for biological replicates of cells re-cultured for 1 and 2 weeks.

### Intracellular concentrations of rucaparib are heterogeneous at the single cell level

Our PDE models suggested that differential drug accumulation is associated with heterogeneity in drug response. To understand whether this was recapitulated in cell line models, single cell measurements of intracellular drug concentrations were required. Rucaparib has intrinsic fluorescent properties that enable its visualisation by fluorescence microscopy^22^, and high-content imaging approaches showed a clear and specific rucaparib signal in PEO1 cells (Extended Data Fig. 6a) that could be visualised as early as 1 minute after addition of the drug, later developing a punctate distribution (Fig. 3c). Quantitation confirmed that intracellular concentrations of rucaparib ([Rucaparib]_IC_) were indeed heterogenous, with a 5-fold difference in signal arising from those cells with the highest versus the lowest drug signal, and a coefficient of variance (CV) of 43.4±9.8% (Fig. 3d). By fitting a one-phase association curve, it was observed that the rate of uptake into cells (K) was inversely correlated to the steady-state [Rucaparib]_IC_ (r^2^ = −0.48±0.12, Extended Data Fig. 6b). Heterogeneity in [Rucaparib]_IC_ was also observed in other OVCA cell lines, with CVs ranging from 43±16% to 77±12% (Fig. 3e).

To understand whether differential rucaparib uptake in PEO1 cells was due to genetic variability within the population, we treated cells with rucaparib for 2 hours (to ensure steady-state [Rucaparib]_IC_ had been achieved, Fig. 3d) and then FACS-sorted the top and bottom 20% of the population, based on rucaparib fluorescence, to obtain ‘Ruc_High_’ and ‘Ruc_Low_’ populations, respectively (Fig. 3f). These populations were then returned to culture for 1, 2 or 3 weeks before re-incubating with rucaparib and analysing the fluorescence distribution once more, to determine whether the phenotype was stable over several rounds of cell division (Extended Data Fig. 6c, Fig. 3f, g). Ruc_High_ and Ruc_Low_ populations had almost identical fluorescence distributions after 3 weeks in culture. Together these data demonstrate that the intracellular concentration of rucaparib is highly heterogeneous at the single cell level, and that this heterogeneity is likely to arise from non-genetically encoded molecular differences between single cells within the bulk population.

### Deconvoluting the molecular drivers of differential PARP inhibitor accumulation from pharmacological consequences

To determine the molecular drivers of differential rucaparib accumulation, we reasoned that by sorting rucaparib-treated cells at early time points, we would enrich for intrinsic molecular differences between the ‘high-’ and ‘low-drug’ populations, while minimising changes driven by differential drug responses (Fig. 4a).

**Figure 4:**
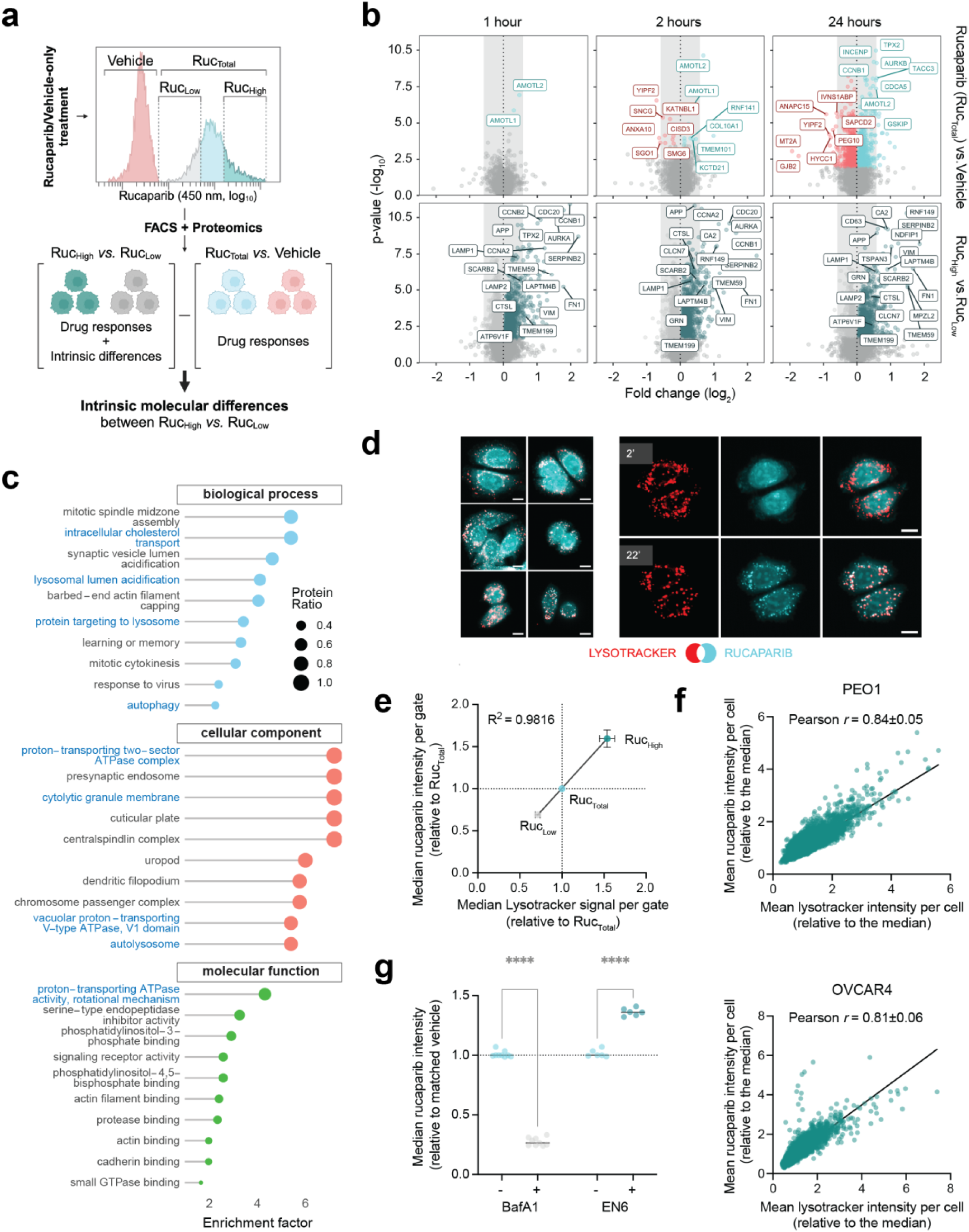
Lysosomal content determines intracellular rucaparib concentration. A) Experimental rationale. Cells are treated with rucaparib or vehicle-only. After treatment, cells are FACS-sorted into Ruc_High_ and Ruc_Low_ populations (top and bottom 20% of the rucaparib positive population), followed by protein extraction, tryptic digestion, and proteomics analysis. Proteins that are differentially expressed between vehicle-only and drug-treated total (Ruc_Total_) populations (control groups) reflect changes in response to intracellular drug concentration. In contrast, proteins differentially expressed between Ruc_High_ and Ruc_Low_ populations—but not significantly altered in control groups—likely represent intrinsic differences pre-existing drug treatment. B) Volcano plots depicting differential protein levels within rucaparib-treated PEO1 cells following a 1, 2 or 24h incubation period. Samples were obtained on 4 different days and 2 technical replicates were collected per day. Proteins with different abundance across conditions (q-value < 0.05) are coloured, with labelling of selected differentially expressed hits. C) Gene ontology pathway enrichment analysis of proteomic differences between our Ruc_High_ and Ruc_Low_ PEO1 populations. Enrichment was calculated for up-regulated proteins at the 1-hour rucaparib treatment time point (q-value < 0.05). A q-value cut-off of 0.05 was applied to define significance of enrichment. Lysosomal-related term are highlighted in blue.D) Left; examples of colocalisation of rucaparib with LysoTracker at 22 minutes post dosing in PEO1 cells. Right; changes in drug distribution at 2 (top) and 22 minutes (bottom) post dosing. Cyan: Rucaparib; Red: LysoTracker Red. 10 µm scale bar. E) FACS-based correlation between lysosomal and rucaparib contents in PEO1 cells, after 1.5h dosing of both compounds. Cell population was gated as before (see section a) and median LysoTracker and rucaparib signal were calculated per gate, relative to the ungated population (Ruc_Total_) per biological replicate (n = 3 technical and biological replicates). F) Correlation between mean lysosomal content per cell (LysoTracker intensity) and mean rucaparib fluorescence intensities in PEO1 (top) and OVCAR4 (bottom) cell lines, after 1hr dosing. Data representative of n = 3 biological replicates. Pearson correlation coefficients were calculated as the mean ± SD across all biological replicates. G) FACS-based quantification of median rucaparib signal in PEO1 cell populations. Cells were pre-treated with BafA1 or EN6 for 1 or 3 hours respectively, followed by 1 further hour with BafA1/EN6 plus rucaparib. Rucaparib signal was normalised to the signal intensity of the vehicle (rucaparib-only) population (BafA1 = 3 biological replicates, EN6 = 2 biological replicates, each biological replicate = 3 technical replicates). Two-way ANOVA and Sidak multiple comparison test were applied to evaluate statistical significance.

A proteomics analysis of the total population of rucaparib versus vehicle-treated cells after just 1 hour of treatment revealed just two significantly changed proteins, suggesting minimal detectable drug response at the protein level at this time point (Fig. 4b). At the same early time point, a large number of proteins were significantly enriched in Ruc_High_ relative to Ruc_Low_ cells, which were inferred to be intrinsic molecular differences between these two populations of cells. Similar expression profiles were also found between Ruc_High_ and Ruc_Low_ cells sorted after 2 and 24h of treatment, suggesting that the proteins that drive differential rucaparib accumulation are maintained as the cells respond to rucaparib treatment (Fig 4b, Supplementary Data Table 6). Pathway enrichment analysis revealed terms that converged strikingly on lysosomal acidification/autophagy (e.g. LAMP1, LAMP2, CTSL, APP, SCARB2, ATP6V1F) (Fig. 4c, Supplementary Data Table 7) recapitulating the findings from our PDE samples. The identification of lysosomal signatures at early time points (where drug responses are minimal) in the Ruc_High_ populations suggests that they are an intrinsic difference between these and the Ruc_Low_ population.

Ruc_High_ cells were also enriched in a number of mitotic proteins (e.g. AURKA, aurora kinase; CCNB1 and CCNB2, cyclin B1 and B2; TPX2). Since G2/M phase cells are larger, we looked at the relative cell sizes in our Ruc_High_ and Ruc_Low_ populations. Though Ruc_High_ cells had a 26% increase in forward-scatter signal (FSC-A, which is proportional to cell diameter), this was not comparable to the 5.7-fold difference in median rucaparib signal, and after normalisation to cell volume, there was still an approximately 2.5-fold difference in median rucaparib fluorescence per cell in Ruc_High_ versus Ruc_Low_ populations at early time points, suggesting that increased cell size in the G2/M phase was not a major driver of the increased drug signal in Ruc_High_, but may instead be a correlating feature with cells that have higher lysosomal content^23^ (Extended Data Fig. 7a-c).

We had noted previously that rucaparib localisation within cells became punctate over time. Given the lysosomal signatures observed in both patient samples and Ruc_High_ cells - and considering that several drugs are known to localise to the lysosome^24–26^– we hypothesised that rucaparib may accumulate there. Live cell imaging of rucaparib with LysoTracker dye demonstrated that, although initial signal is cytosolic, rucaparib punctae colocalise with lysosomes within 30 minutes of drug incubation (Fig. 4d, Extended Data Fig. 7d). Cells treated with both rucaparib and LysoTracker displayed a striking correlation between both fluorescent signals, with Ruc_High_ being significantly enriched in LysoTracker signal with respect to Ruc_Low_ (Fig. 4e, Extended Data Fig. 7e, Pearsons rho = 0.83±0.08). Image quantification in PEO1 and OVCAR4 cells also revealed that LysoTracker signal was highly correlated with rucaparib levels per cell (Pearson rho = 0.84±0.05 and 0.81±0.06 respectively; Fig. 4f). Transient over-expression of TFEB-GFP, a master regulator of lysosomal biogenesis, significantly increased [Rucaparib]_IC_ demonstrating that lysosomal content changes can drive increased rucaparib accumulation (Extended Data Fig. 7f, g). Conversely, treatment of cells with bafilomycin or chloroquine (CQ) to alkalinise lysosomes decreased [Rucaparib]_IC_ (Fig. 4g, Extended Data Fig. 7h-j). Additionally, the V-ATPase activator EN6^29^ triggered an increase in intracellular rucaparib levels (Fig. 4g). Together, these findings demonstrate that heterogeneous [Rucaparib]_IC_ is driven by differential lysosomal content, and that this accumulation is pH-dependent.

### Differences in intracellular Rucaparib concentration correlate with DNA damage response in equimolar treated cells

To determine whether differential intracellular concentrations of PARP inhibitors could be a contributing factor to the variable DNA damage response seen in PEO1 and other OVCA cell lines, measurement of [Rucaparib]_IC_ and ɣH2AX levels were required within the same cell.

PARP inhibitor-induced DSBs occur as a result of PARP trapping and replication fork collapse during S-phase^14^. We therefore synchronised cells in G1 phase with palbociclib^27,28^ (Extended Data Fig. 8a, b) prior to releasing them in the presence of rucaparib, to ensure that all cells that had reached G2/M after treatment had had equal opportunity to accumulate DNA damage. We found that indeed, in palbociclib-synchronised cells, there was a positive correlation between [Rucaparib]_IC_ and ɣH2AX levels, and that this correlation was strongest in cells in G2/M (Fig. 5b, c), in keeping with this population having the largest ‘opportunity’ to accumulate damage. Furthermore, separation of cells into quartiles based on levels of intracellular rucaparib, revealed a statistically significant increase in ɣH2AX from one quartile to the next (Fig. 5d) with a linear relationship between the median [Rucaparib]_IC_ and median ɣH2AX fluorescence intensity within each quartile (Fig. 5e). This interaction was also observed in OVCAR4 cells (Extended Data Fig. 8c, d).

**Figure 5:**
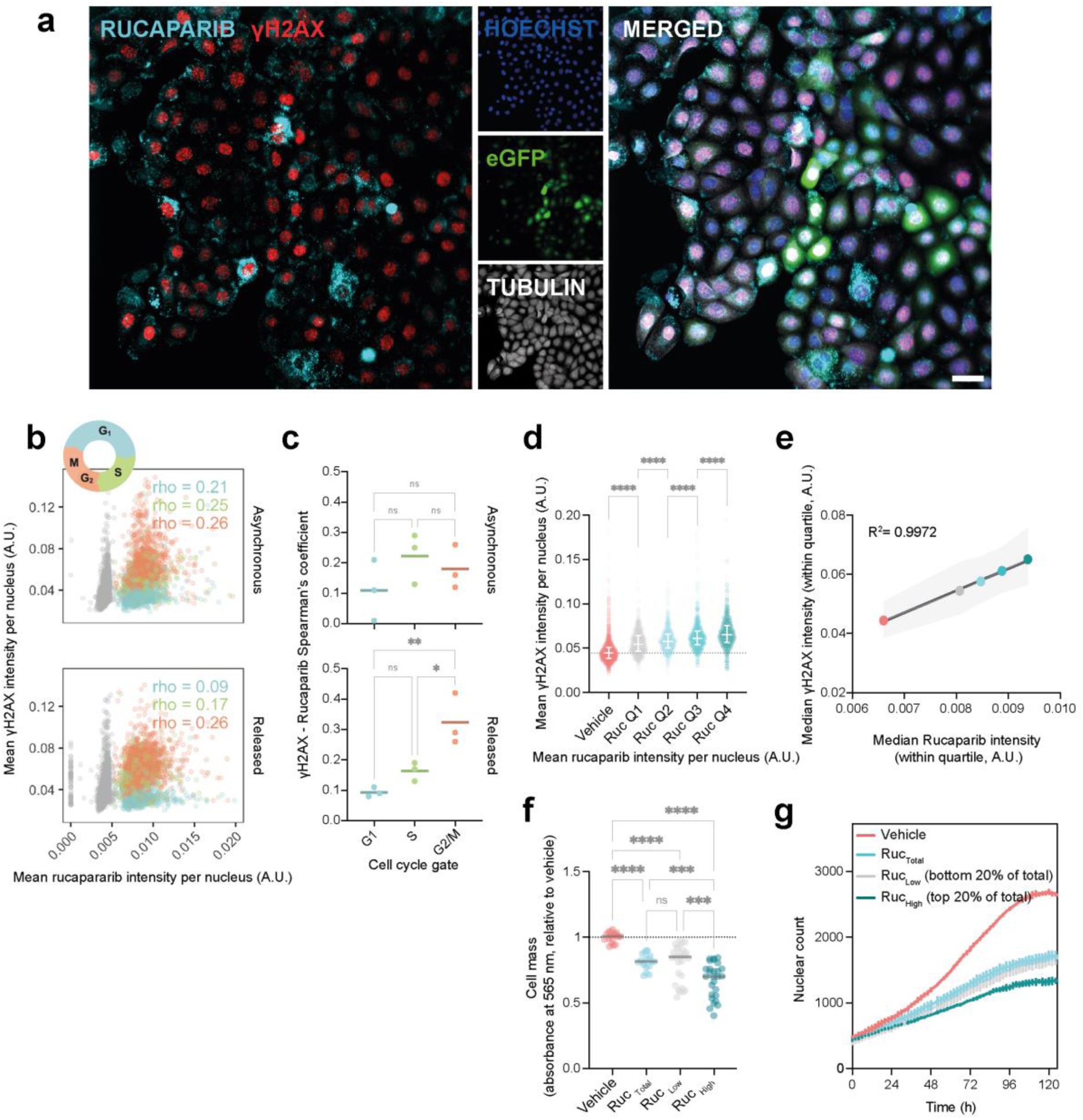
Differences in intracellular rucaparib concentration contribute to heterogeneity in DNA damage levels in equimolar-treated cells. A) Rucaparib and γH2AX imaging in PEO1 cells. Rucaparib (355/480 nm) and eGFP (489/511 nm) were imaged live. Cells were fixed and immunostained for γH2AX and tubulin, using hoechst to stain the nuclei. eGFP was imaged a second time, to use as a common element to co-register live and fixed-cell images. Scale bar = 50 μm. B) Palbociclib-treated (‘released’) and asynchronous PEO1 cells were treated with rucaparib at IC_50_ concentration or vehicle-only for 24 hours. Cells were then imaged following the procedure described in A) to correlate rucaparib and γH2AX levels per nuclei. Spearman’s correlation coefficients across cell cycle gates (blue: G1 phase; green: S phase; red: G2 phase) indicated for both asynchronous control and palbociclib-released conditions. Greyed data points represent the vehicle-only population. p < 0.0001 for all *rho* values except released-G1 condition, with p = 0.0176 (n=3 technical replicates). Data representative of n=3 biological replicates. C) Cell-cycle resolved Spearman’s correlation coefficients between rucaparib and γH2AX levels per biological replicate (n = 3). One-way ANOVA with Tukey’s multiple comparison tests were used. D) The G2/M gate of the palbociclib-treated PEO1 cell population was subdivided into quartiles based on nuclear rucaparib signal to compare γH2AX levels between groups. Data representative of n = 3 biological replicates. One-way ANOVA with Tukey’s multiple comparison tests were used. E) Median γH2AX is linearly related to median intracellular rucaparib within the quartile. Pooled data from n = 3 biological replicates is depicted. F) Following a 2-hour treatment, PEO1 cells were FACS-sorted into Ruc_High_ and Ruc_Low_ populations (top and bottom 20% of the rucaparib positive population). Cells were re-cultured, allowed to proliferate in drug-free medium for 72 hours, and stained with SRB to measure cell mass. Data representative of n = 5 biological replicates, with n = 3 technical replicates, is depicted. One-way ANOVA with Tukey’s multiple comparison tests with a single pooled variance were used to assess statistical significance. G) Growth curves depicting re-proliferation of the different gates following rucaparib-FACS sorting. Data shows n = 3 technical replicates and is representative of n = 3 biological replicates. Differences in growth rates and plateaus were confirmed in an extra sum-of-squares F test over the fitted logistic growth model, with p < 0.0001.

To understand the implications of differential rucaparib accumulation on drug response, cells were again sorted into Ruc_High_ or Ruc_Low_ populations. Even after just 2 hours of treatment, Ruc_High_ cells subsequently proliferated more slowly and plateaued at a significantly lower cell mass than Ruc_Low_ cells (Fig. 5f, g), further demonstrating that differential uptake at the single cell level impacts rucaparib response. Together, these data demonstrates that despite equimolar dosing under standard cell culture conditions, differences in [Rucaparib]_IC_ contribute to heterogeneity in DNA damage and rucaparib efficacy at the single cell level. Since lysosomal accumulation is highly correlated to intracellular rucaparib levels, which, in turn, are associated with increased DNA damage, this suggests that lysosomal accumulation of rucaparib increases drug efficacy.

#### Lysosomal accumulation increases nuclear bioavailability of weak base PARP inhibitors

Lysosomal localisation of drugs is often referred to as trapping, and therefore associated with drug resistance. However, Ruc_High_ cells have increased lysosomal accumulation and yet have increased drug response. We therefore hypothesised that lysosomes act as a reservoir that maintains [Rucaparib]_IC_ at a higher level and therefore improves bioavailability. Since PARP1 and 2 are nuclear proteins, we quantified nuclear rucaparib per cell, with or without the addition of bafilomycin, CQ, or the V-ATPase activator EN6 (Fig. 6a, Extended data Fig. 9c, h). Alkalinisation of lysosomal pH with bafilomycin or CQ treatment prevented lysosomal accumulation and substantially decreased nuclear levels of rucaparib in PEO1 cells (Fig. 6a, b, Extended data Fig. 9a-e). Similar results were seen for OVCAR4 cells treated with bafilomycin (Extended data Fig. 9h-j). Conversely, increased acidification of lysosomes with EN6 drove increased lysosomal accumulation and elevated nuclear rucaparib levels in PEO1 cells (Fig. 6b, Extended data Fig. 9f, g). Upon washout of rucaparib from the extracellular space, rate of fluorescence loss was slower in cells with intact lysosomal function, relative to those treated with bafilomycin or CQ, further supporting the model that the equilibrium of cytosolic and nuclear rucaparib is maintained at a higher concentration by lysosomal accumulation (Fig. 6c, Extended Data Fig. 9k).

**Figure 6:**
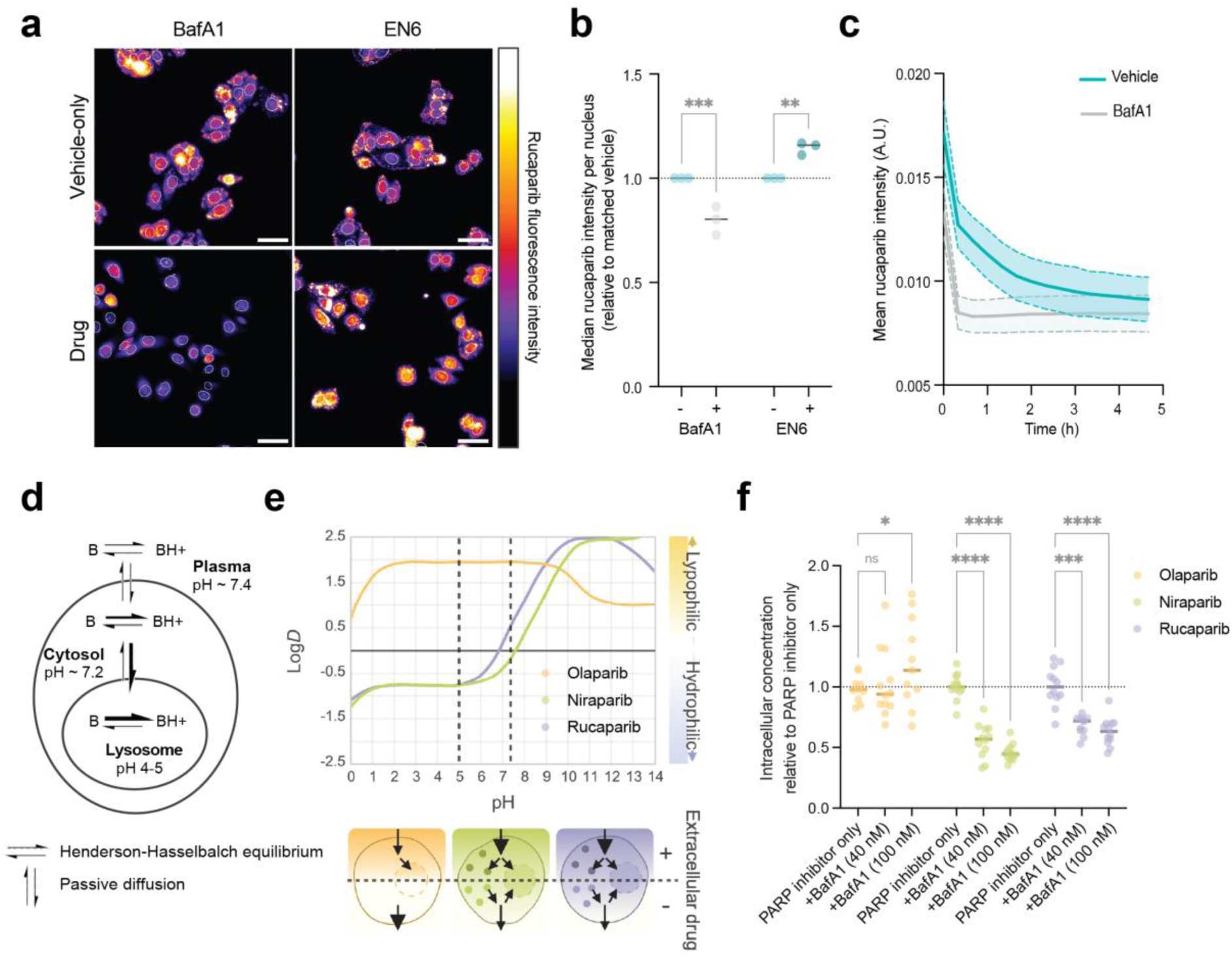
Lysosomal localisation increases the nuclear bioavailability of niraparib and rucaparib but not olaparib. A) Representative images of rucaparib in PEO1 cells after 1 hour with or without pre-treatment with the V-ATPase inhibitor bafilomycin (BafA1) or the V-ATPase activator EN6. Dotted white lines outline the nucleus. Scale bar = 50 µm. B) Imaging-based quantification of nuclear levels of rucaparib in PEO1 cell populations pre-treated with BafA1 or EN6 for 1 hour, followed by 1 further hour with BafA1/EN6 plus rucaparib. Rucaparib signal was normalised to the signal intensity of the vehicle (rucaparib-only) population (Data shown from n = 3 biological replicates, each point shown is mean of n = 3 technical replicates). Two-way ANOVA and Sidak multiple comparison test were applied to evaluate statistical significance. C) Rucaparib fluorescence loss from PEO1 cells over time after rucaparib is removed from the medium, with or without bafilomycin. Data shows mean rucaparib fluorescence intensity per cell ± standard deviation (shaded area) calculated across 6 replicate wells and is representative of 2 independent biological replicates. D) Diagram to represent pH-dependent partitioning of lipophilic, weak basic drugs. Adapted from^30^. E) Top; Illustration of changes in LogD of olaparib, rucaparib and niraparib at different pH, demonstrating the large shift in lipophilicity of rucaparib and niraparib at lysosomal pH (< 5) relative to cytosolic pH (7.2). Bottom; schematic illustrating lysosomal accumulation driving increased intracellular concentrations of niraparib and rucaparib and sustained target engagement under drug-washout conditions. F) Intracellular concentration of olaparib, niraparib and rucaparib in PEO1 cells with or without treatment with bafilomycin to alkalinise lysosomes. Data represents combined data from 3 biological replicates, each within 4 technical replicates, and plotted relative to PARP inhibitor only control, within biological replicate. Statistical significance assessed by 2-way ANOVA with Dunnet’s correction for multiple comparisons.

Our PDE spatial transcriptomics data suggested an interaction with lysosomes for niraparib as well as rucaparib. Lysosomal accumulation has been documented for a range of drugs^26,30–34^, particularly those that are weak bases. These compounds become protonated in the acidic environment of the lysosomal lumen, a process that reduces their lipophilicity (LogD) and their ability to diffuse back into the cytosol—ultimately leading to their accumulation within the organelle (Fig. 6e). In the case of rucaparib, our data suggest that an equilibrium is maintained between the lysosomal pool of drug and the cytosol/nucleus, with lysosomal accumulation thereby maintaining higher drug concentrations across the cell. Among the PARP inhibitors clinically used, both rucaparib and niraparib are weak bases, with p*K*_a_s of 9.3 and 10.1 for their most basic moieties respectively (drugbank), thus making them susceptible to this mechanism. Conversely, olaparib has a much lower p*K*_a_ (−0.9), suggesting it is unlikely to undergo pH-dependent lysosomal trapping (Fig. 6f). To test this, as neither niraparib nor olaparib are fluorescent, we measured intracellular concentrations of all three PARP inhibitors, with or without bafilomycin pre-treatment, using liquid chromatography-mass spectrometry. Both rucaparib and niraparib intracellular concentrations were significantly decreased in PEO1 and OVCAR4 cells in the presence of bafilomycin, whereas olaparib was unaffected (Fig. 6g, Extended Data Fig. 9l), supporting the hypothesis that only weak base PARP inhibitors accumulate lysosomally.

Together, these data suggest that as weak bases, rucaparib and niraparib accumulate within the lysosome and that this accumulation increases their bioavailability, whereas olaparib bioavailability is unaffected by lysosomal content or function.

## Discussion

The intracellular concentration of a drug is central to determining its target binding and efficacy: too low, and the intended target will not be engaged; too high, and the drug will interact with lower affinity interactors, leading to off-target effects^15^. Despite its importance, drug distribution within tumours, and the mechanisms that regulate this, are poorly understood. Following equimolar *ex vivo* dosing in a vasculature-independent, patient-derived system, our novel multi-modal imaging pipeline demonstrated the presence of heterogeneity of PARP inhibitor distribution at three levels: between patients, different tumour sites from the same patient, and within individual tumours.

Interestingly, while niraparib and rucaparib exhibited similar relative uptake levels in a given patient, olaparib followed a distinct pattern, suggesting that the mechanisms of accumulation differ between olaparib and the other two drugs. This observation was borne out by the fact that intracellular concentrations of niraparib and rucaparib are decreased by lysosomal alkalinisation, however olaparib is not.

It has been shown that a number of drugs accumulate within the lysosome^26,35^, with the suggestion that this occurs because they are weak bases that become protonated within the acidic lysosomal lumen. Lysosomal sequestration is often referred to as “drug trapping” which limits cytosolic or nuclear target engagement, but this mechanism of resistance remains poorly characterised. Furthermore, the term “trapping” can be misleading, as weakly basic drugs accumulated in lysosomes may still diffuse back into the cytoplasm under a favourable concentration gradient^30^. Our experiments suggest that the lysosomal pool of rucaparib and niraparib remains active and enhances drug function. This is evidenced by increased DNA damage in Ruc_High_ cells, and decreased nuclear levels of rucaparib when lysosomal accumulation is diminished through bafilomycin or chloroquine pre-treatment. This fits a model where lysosomes act as a reservoir of rucaparib or niraparib, enabling the cell to maintain an overall higher concentration at the target site, analogous to the accumulation of the tuberculosis drug bedaquiline in lipid droplets^36^. In line with this, we see that rucaparib concentrations are maintained for longer in cells with an intact lysosomal compartment. Our findings highlight the complexity of the role of lysosomes in therapeutic sensitivity and resistance of PARP inhibitors, and other weak base drugs, and warrants further investigation.

The work presented here is of clinical importance: regions of high niraparib and rucaparib accumulation in PDEs displayed increased apoptotic signatures, while Ruc_High_ PEO1 cells had decreased proliferation rates after just two hours of exposure, suggesting better drug responses in higher drug accumulating regions, potentially enabling lower-drug accumulating regions to drive therapeutic resistance. Furthermore, since tumours used here were resected from chemotherapy-naïve patients, our work highlights that differential accumulation of PARP inhibitors may be a contributing factor to intrinsic resistance to these drugs. Clinically, PARP inhibitors are dosed orally and will reach the tumour via the vasculature, whose often disorganised structure is likely to further increase the heterogeneity of distribution of these drugs across the tumour.

In summary, our PDE and cell-based studies reveal that lysosomal accumulation of rucaparib and niraparib can influence drug efficacy, while olaparib appears to accumulate through a lysosome-independent mechanism. This difference could suggest that if poor pharmacodynamics drives lack of response to one PARP inhibitor within a patient’s tumour, treatment with another may still be effective. Our multi-modal imaging study also opens the possibility of personalised therapy, where biopsy samples can be used to assess drug accumulation, and to determine which drug is likely to be most effective for a patient. Finally, our findings highlight cell-intrinsic variability in drug accumulation within patient samples. This clinically relevant evidence underscores the need to investigate how tumour heterogeneity affects drug distribution as a potential mechanism of intrinsic resistance across other cancer therapies and tumour types.

## Methods

### Spatially resolved approaches using patient-derived tumour explants

Human HGSOC tumour samples were obtained through intraoperative tumour mapping^37,38^ from chemo-naive patients under-going maximal effort upfront cytoreductive surgery at Hammersmith Hospital, London, UK, a tertiary gynaecological cancer centre, certified by European Society of Gynaecological Oncology (ESGO) as centre of excellence for ovarian cancer surgery^39,40^. The human samples for this research project were banked by the Imperial College Healthcare Tissue Bank (ICHTB). ICHTB is supported by the National Institute for Health Research (NIHR) Biomedical Research Centre based at Imperial College Healthcare NHS Trust and Imperial College London. ICHTB is approved by Wales REC3 to release human material for research (22/WA/2836). All patients gave written consent. The procedures involved human participants were done in accordance with the ethical standards of the institutional and/or national research committee and with the principles of the 1964 Declaration of Helsinki and its later amendments or comparable ethical standards.

#### Tissue slicing and culture

Prior to processing, samples were kept on ice. PDEs were obtained using a vibrating blade microtome (LeicaVT1200 S) to cut slices of 400 to 600 μm in thickness. 5% agarose in PBS was used to embed tumour samples prior to slicing, which was performed in PBS supplemented with 100 U/mL penicillin, 100 μg/mL streptomycin (P/S). Resulting PDEs were placed onto inserts (0.4 μm porosity, Millipore PICM01250) within 6-well plates and just submerged for acclimatisation overnight in complete medium (RPMI 1640, 20% FCS (foetal calf serum), 2 mM sodium pyruvate, 100 U/mL penicillin, 100 μg/mL streptomycin, 2.5 μg/mL insulin) at 37°C, 5% CO_2_.

#### Ex vivo dosing with PARP inhibitors

Explants were treated with niraparib, olaparib and rucaparib at 20 μM in complete medium or with 0.1% DMSO as vehicle. PARP inhibitors have a C_max_ (source: drugbank) in the micromolar range (12.4-17.5 µM, 6 µM and 2.5 µM for olaparib, rucaparib and niraparib respectively), therefore the concentrations used for explants are in line with those observed in patients. PDEs were incubated with drug (37°C, 5% CO_2_) for between 2 and 48 hours. Typically, 24 hours treatment was used to ensure homogeneous drug distribution throughout the tissue slice. Post-incubation, ice-cold drug-free medium was used to rapidly wash PDEs before rinsing with ice-cold PBS. Excess liquid was carefully blotted before snap-freezing in liquid nitrogen. Samples were stored at −80°C.

#### Embedding and cryosectioning

To aid cryosection, frozen PDEs were embedded in a hydrogel (7.5% hydroxypropyl methylcellulose, 2.5% PVP)^41^. Polymer blocks were snap-frozen using dry ice-chilled isopropyl alcohol. 10-12 μm serial sections were cut using a Leica CM3050s cryostat (−17°C), and thaw-mounted onto slides (Thermo Fisher, 12-550-15) prior to storage at −80°C.

#### Histochemistry and immunohistochemistry

Acetone was used to fix sections (−20°C). H&E staining was undertaken using the commercial kit ab245880 (Abcam) according to manufacturer’s instructions, before mounting with DPX. For IHC, endogenous peroxidase activity was quenched using 3% hydrogen peroxide. Sections were then incubated with protein blocking buffer (Abcam) for 1 hour at room temperature, followed by overnight incubation at 4°C with primary antibodies diluted in either SignalStain Antibody Diluent or the same blocking buffer. Slices were then incubated with secondary antibodies for 1 hour (anti-rabbit IgG conjugated with polymeric horseradish peroxidase linker (Leica Bond Polymer Refine Detection, DS9800) for γH2AX and cleaved caspase-3 primaries; biotinylated goat anti-rabbit IgG (HRP/DAB detection IHC Kit, ab64261) for the rest). DAB was used as chromogen and sections were haematoxylin counterstained and mounted with DPX. Slides were imaged using a ZEISS Axioscan Z1 slide scanner and a DM4b/DM6000 upright Leica setup and processed with Fiji and QuPath^42,43^.

#### Mass spectrometry imaging (MSI)

Tissue sections were thawed under low vacuum for 20-30 minutes. For method development and standard curve calculation, 0.15 μL of compound of interest at known concencentrations (diluted in 50% MeOH, 1% DMSO as vehicle) were spotted onto desiccated tissue sections, before further processing. Two MSI platforms were used as described below. Initial set up of our multimodal imaging pipeline (Extended Data Fig. 1e, f) used DESI-MSI which has a lower spatial resolution but improved speed of acquisition. All MSI completed as part of the main study (Fig. 1, Extended Data Fig. 2) used the MALDI MSI platform to maximise spatial resolution.

#### Matrix-assisted laser desorption/ionization (MALDI) MSI

Matrix (2,5-Dihydroxybenzoic acid (DHB, 20 mg/mL in 50% acetone, 0.1% TFA)) was applied to the tissue section which was held by a rotating target (350 rpm) at 5 μL/min (nitrogen flow rate of 5 L/min) for 30 minutes using a pneumatic sprayer (SMALDI prep, TransMIT GmbH). Heavy-isotope labelled versions of each of the PARP inhibitors ([^13^C_6_]-Niraparib, [^2^H_8_]-Olaparib and [^13^C,^2^H_3_]-Rucaparib; Aslachim) were spiked into the matrix solution at 3 μg/mL as internal standards (IS) as described elsewhere^44^. Positive ion mode was used to acquire spectral data, typically between 250-1000 m/*z*, using an AP-SMALDI5-AF ion source (TransMIT GmbH) coupled to a Q-Exactive Plus mass spectrometer (Thermo Fisher Scientific). Source conditions: capillary temp 350-400 °C; voltage, 4 kV. Mass spectrometric parameters: inject time, 250 ms; 70,000 mass resolution; 3.8 scans/s. Scanning was performed in 2D-line mode with a nominal pixel size set to 20 μm, and laser attenuator set to 34 arbitrary units.

After acquisition, cold ethanol was used to remove matrix to enable H&E staining of the tissue section. Raw spectral data was converted to imzML format (RAW2IMZML converter, TransMIT GmbH). Mirion (TransMIT JLU) and MSiReader^45^ were used for data analysis. Normalisation to the appropriate heavy labelled IS was performed. A tolerance window of 5-10 parts per million (ppm) was used to plot features of interest.

#### Desorption Electrospray Ionization (DESI) MSI

For DESI imaging a source comprising a 2D sample holder moving stage and custom-built inlet capillary (490°C) (Prosolia, Indianapolis, USA), coupled to a XEVO G2-XS QToF (Waters Corporation, Milford, MA) was used. DESI parameters were as follows: spray voltage, 4.5 kV; solvent, methanol/water, 95:5; flow rate, 2 μL/min; nebulizing gas, nitrogen; gas pressure, 5 bar; sprayer incidence angle, 75°; collection angle, 10°; sprayer-to-inlet distance, 1 mm; sprayer-to-sample distance, 1 mm. Mass spectrometric parameters: source temperature, 150 °C; source offset −80 V; mass resolution 20,000; 1 scans/s. A nominal pixel size of 75 μm was used. Spectral data was acquired in both positive and negative ion modes between m/z 50 to 1000. Images were generated using HDImaging v.1.4 and MassLynx v4.1 software (Waters Corporation, Milford, MA), applying total ion count (TIC) normalisation, with m/z tolerance value of 20 ppm.

#### GeoMx DSP spatial transcriptomics

Adjacent tissue sections to those used for MSI were processed as recommended by Nanostring. Briefly, samples were thawed, fixed overnight in 10% neutral buffered formalin (NBF), and washed in PBS before baking for 30 mins at 60°C. They were then dehydrated with successive washes (5 minutes) in increasing ethanol concentrations. Antigen retrieval was performed at 100°C for 15 minutes in Tris-EDTA. Proteinase K was used at 1 μg/mL in PBS to expose RNA targets (37°C, 15 minutes). Post-fixation step followed, samples were incubated for 5-minutes in 10% NBF and washed twice in NBF stop buffer (101 mM Tris, 100 mM Glycine in DEPC treated water). In situ hybridization using Nanostring’s Whole Transcriptome Atlas probe mix (overnight, 37°C), was followed by two 25-minute incubations in stringent wash (50% deionized formamide, 2X SSC buffer, 37°C), to remove off-target probes. Samples were blocked with buffer W (30 minutes, NanoString). Finally, hybridised tissue samples were stained with the nucleic acid dye SYTO-13 and the fluorophore-conjugated, primary antibodies (1 hour at room temperature) detailed in Table 2. Hybridised and stained samples were scanned on the GeoMx DSP platform, and regions of interest (ROI) selected, based on overlaid MALDI-MSI images of PARP inhibitor distribution in adjacent tissue sections and morphology marker staining. Typically, a minimum of 80 nuclei were selected for each ROI to ensure sufficient sequencing coverage. PCR amplification/library preparations was carried out according to manufacturer’s instructions. Pooled libraries were purified using AMPure XP beads (Beckman Coulter), and final library quality and quantity assessed using the Agilent 2100 High Sensitivity DNA, and Qubit High-Sensitivity DNA assays.

**Table 1:** Immunohistochemistry primary antibodies.

| Target | Antibody Clone | Species | Dilution |
| --- | --- | --- | --- |
| Wilms Tumour Protein (WT1) | Abcam, ab89901 CANR9(IHC)-56-2 | Rabbit | 1:500 |
| Pax8 | Abcam, ab189249 EPR13511 | Rabbit | 1:1000 |
| Phospho histone H2A.X (Ser139) | CST, 2577 | Rabbit | 1:400 |
| Cleaved Caspase-3 (Asp175) | CST, 9664 5A1E | Rabbit | 1:100 |

**Table 2:**
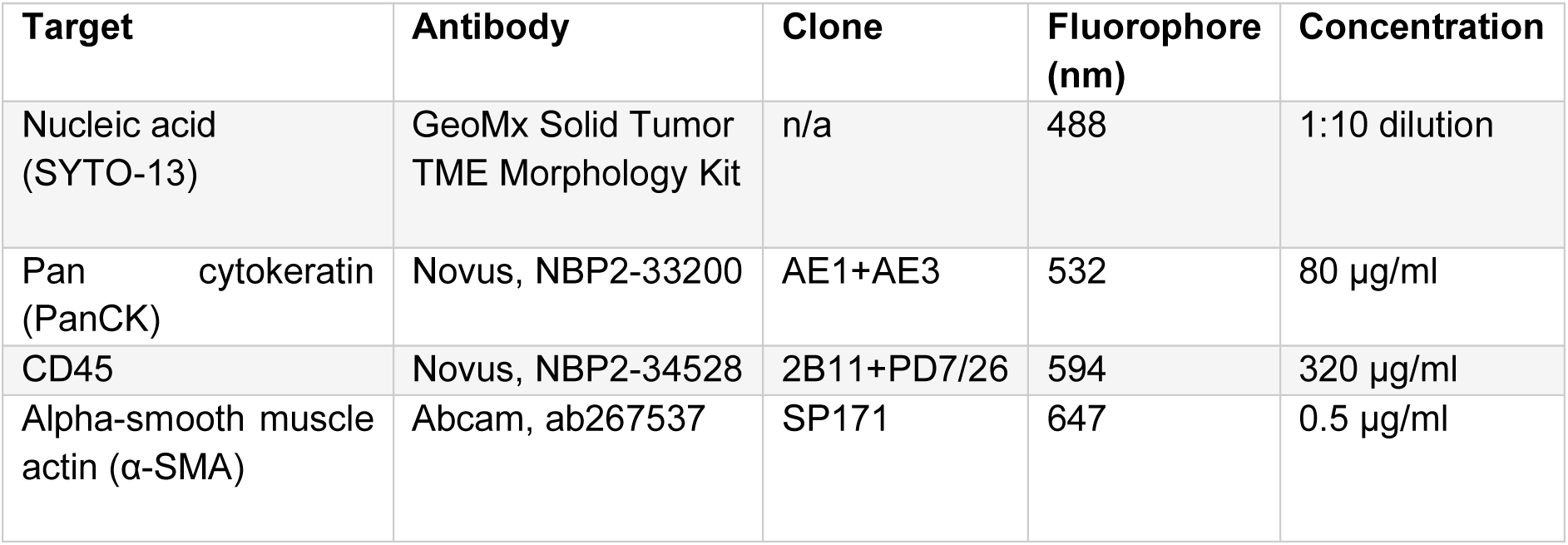
Spatial Transcriptomics staining reagents.

| Target | Antibody | Clone | Fluorophore (nm) | Concentration |
| --- | --- | --- | --- | --- |
| Nucleic acid (SYTO-13) | GeoMx Solid Tumor TME Morphology Kit | n/a | 488 | 1:10 dilution |
| Pan cytokeratin (PanCK) | Novus, NBP2-33200 | AE1+AE3 | 532 | 80 µg/ml |
| CD45 | Novus, NBP2-34528 | 2B11+PD7/26 | 594 | 320 µg/ml |
| Alpha-smooth muscle actin (α-SMA) | Abcam, ab267537 | SP171 | 647 | 0.5 µg/ml |

Sequencing was performed using the Illumina NextSeq2000 platform (Paired End 27bp + dual 8bp indexing). Sequenced data were analysed using the GeoMx NGS pipeline. Quality control thresholds: 100,000 raw reads and 50% sequencing saturation. Quartile3 normalisation was applied across all targets to account for differing ROI size and cellularity which impact transcript abundance. ROIs with 5% or more targets above the limit of quantification (LOQ, defined as negative probe geomean x negative probes geometric standard deviation) were further processed. Targets identified above LOQ greater than 10% of ROIs were included in the analysis. Genes that were differentially expressed in ‘high-drug’ relative to ‘low-drug’ ROIs were identified using a linear mixed model (LMM), using ‘Patient ID’ and ‘PDE slice ID’ as correction variables. Benjamini and Hochberg False Discovery Rate multiple test correction was applied, with a significance threshold of q < 0.05. Gene Set Enrichment Analysis^46^ was carried out using Reactome^47^.

#### Modelling spatial transcriptomics data using drug concentration as a continuous variable

Following quality control and Q3 normalisation, we used the R package *lme4* for linear mixed model analysis of relationship between local drug concentration and gene expression. To model drug concentration (DIC) as a continuous variable (rather than binary ‘high’ versus ‘low), signal intensities corresponding to each ROI were quantified from MSI images, calculating the mean signal intensity within each ROI, relative to heavy-labelled drug-analogue internal standards. To address the hierarchical structure of the data, we employed a linear mixed model with patient and PDE slice as nested random effects, accounting for both inter-patient variability and intra-tumour variability across different slices. The nesting structure recognises the likelihood that slices from the same patient will exhibit more similarity than slices from different patients. The model is represented as:

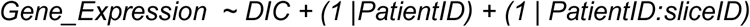

This model was applied to each drug:gene combination separately, and allowed random intercepts at both patient and slice levels, capturing the inherent variability at each level of the hierarchy. To assess the significance of drug effects on gene expression, the P-value was calculated by comparing the full model (including DIC as a predictor) with the null model (excluding DIC), using the R anova() function:

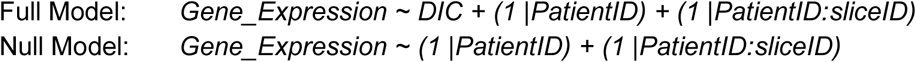

Functional enrichment analysis was performed using Gene Ontology (GO) terms and Reactome pathways. GO term enrichment analysis used the *gseGO* function in the *clusterProfiler* package. Pathway enrichment was analysed using the *gsePathway* function of the *ReactomePA* package.

### General cell-based methods

#### Cell line handling

PEO1 and 4 were a gift from Simon Langdon. ES2, OVCAR8, OVCAR4 and Kuramochi lines were all obtained from ATCC. OVCAR4 EGFP-PARP1-mCherryFP cell line was generated previously^48^. All cell lines were recently authenticated using STR profiling. Cell lines were maintained in RPMI 1640 supplemented with 2 mM Glutamine, 100 U/mL penicillin, 100 µg/mL streptomycin, 10% FCS at 37°C, 5% CO_2_ and passaged three times per week using TrypLE express.

#### Drug dosing

Table 3 details drug used throughout the study. Stock aliquots in DMSO were stored at −80°C. For use, drug stocks were diluted in medium, with a maximum vehicle concentration of 0.1%. Chloroquine stocks were made fresh immediately before use.

**Table 3:** Drug stock details.

| Compound | Catalogue number | Stock concentration | Vehicle |
| --- | --- | --- | --- |
| Bafilomycin A1 | Sigma 5.08409 | 0.1mg/mL | DMSO |
| Chloroquine | Sigma C6628 | 50 mg/mL | PBS |
| EN6 | MCE HY-128892 | 50 mM | DMSO |
| Niraparib | Selleck S2741 | 100 mM | DMSO |
| Olaparib | Selleck S1060 | 100 mM | DMSO |
| Palbociclib | ApexBio A8316-APE | 1 mM | DMSO |
| Puromycin | Merck 540222 | 1mg/mL | H2O |
| Rucaparib | ApexBio A4156-APE | 100 mM | DMSO |

#### Sulforhodamine-B cell mass accumulation assay

For the estimation of drug IC_50_ values, cells were plated in 96-well plates (Corning, 3596) at 4000-6000 cells/well, (cell line-dependent) in complete medium and left to adhere overnight. Cells were dosed as indicated and after 72 hours, plates were fixed with 10/% trichloroacetic acid (1 hour, 4°C) and stained with 0.4% Sulforhodamine-B (30 minutes room temperature). Plates were washed (1% acetic acid) and air-dried before solubilising the stain (200 µL, 10 mM Tris-Base) and reading optical density at 565 nm.

#### Growth curves

Cells were seeded on a clear bottom, white 96-well plate (Corning, 3610) in media, ± 1:1000 dilution of IncuCyte NucLight Rapid Red Reagent (Sartorius) to label nuclei, and cultured at 37°C, 5% CO_2_ while imaging every 2-hours in an Incucyte S3 (Sartorius). Confluency and/or nuclear number was plotted.

#### Immunofluorescence and high content analysis microscopy

Cells were seeded at approximately 40% confluency on 96-well plates (clear bottom, black, Phenoplates, Perkin Elmer, 6055300), and the following day, treated with drug (concentrations and timings indicated in text or legends), before fixing with 4% paraformaldehyde (20 minutes at room temperature). Blocking and permeabilization were combined using 2% bovine serum albumin (BSA), 0.5%Triton-X in PBS for 30 minutes. Primary antibodies in Table 4 were incubated overnight at 4°C at the specified concentration in 2% BSA. Fluorophore-conjugated secondary antibodies (Table 5) were incubated in the dark, typically at a 1:1000 dilution, for 1 hour at room temperature. Hoechst 33342 DNA staining was performed prior to imaging (1 µg/mL in PBS, 15 minutes). Images acquisition and analysis was carried out using the Operetta CLS High Content Analysis System, using the 20x air NA 0.8 objective unless otherwise stated. Five images per well of a single focal plane were acquired using a random pattern while excluding the well borders. Focus was manually set. Harmony Software (Revvity, Inc.) was used for flatfield correction and nuclear segmentation was based on hoechst staining. Measurements such as hoechst sum intensity per nucleus (cell cycle gating), as well as γH2AX fluorescence intensity were recorded and downstream analysis used custom scripts in RStudio. Further additional image processing used Fiji^42^.

**Table 4:** Immunocytochemistry primary antibodies.

| Target | Antibody Clone | Species | Dilution |
| --- | --- | --- | --- |
| Phospho histone H2A.X (Ser139) | CST 2577 | Rabbit | 1:1000 |
| α-Tubulin Sigma | T9026 DM1A | Mouse | 1:1000 |

**Table 5:** Immunocytochemistry secondary antibodies.

| Antibody | Species | Invitrogen catalogue number |
| --- | --- | --- |
| Alexa Fluor 488 | Rabbit | A21206 |
| Alexa Fluor 555 | Rabbit | A31572 |
| Alexa Fluor 568 | Mouse | A10037 |
| Alexa Fluor 647 | Mouse | A31571 |

#### High-content microscopy of rucaparib

Rucaparib was imaged in live cells that had been washed in FBS-containing medium post treatment immediately before imaging (using the hoechst channel in Table 6), with the Operetta CLS High Content Analysis System (set at 37°C, 5% CO_2_), using the 20x air NA 0.8 objective unless otherwise stated. In some experiments, cells were pre-incubated at 37°C with LysoTrackerTM Deep Red (50 nM, 30 mins, Invitrogen L12492, imaged using the Alexa Fluor 647 channel in Table 6) and/or CellTracker Green (10 µM, 15 mins, Invitrogen C7025, imaged using the Alexa Fluor 488 channel in Table 6) and pre-treated with CQ, bafilomycin A1 or EN6 (typically at 25 µM, 50-100 nM and 50 µM respectively) for 1 hour before rucaparib was added for a further hour. When needed, samples were fixed and immunostained following live imaging (as above). Custom Fiji macros were developed to align the resulting images based on the identification of common features from both image sets. Typically, eGFP-overexpressing (see lentiviral transduction below) or CellTracker stained cells were used to enable the initial live and then fixed 488 signal for co-registration. Pearson correlation coefficients between the superimposed 488 channels were calculated pre- and post-alignment to QC the registration process. Should the post-alignment coefficient be lower than the original, the image set was flagged and excluded from analysis. CellProfilerTM 4.2.6 software^49^ was used to analyse aligned images. α-tubulin and hoechst staining was used to segment individual cells and nuclei, respectively. Cytoplasmic segmentation was achieved by subtracting nuclear space from individual segmented cells. Rucaparib fluorescence intensity was quantified for each segment (cell, cytoplasm, nucleus), while γH2AX and hoechst fluorescence intensities were recorded for each nucleus. Downstream analysis was carried out using custom scripts in RStudio.

**Table 6:** Operetta CLS High Content Analysis System channels.

| Channel | Excitation wavelength range (nm) | Emission wavelength range (nm) |
| --- | --- | --- |
| Alexa Fluor 488 | 460-490 | 500-550 |
| Alexa Fluor 555 | 530-560 | 510-650 |
| Alexa Fluor 568 | 530-560 | 510-650 |
| Alexa Fluor 647 | 615-645 | 655-760 |
| Hoechst | 360-400 | 410-480 |
| eGFP | 460-490 | 500-550 |

**Table 7:** Expression plasmids and details for viral vector production.

| Name | Backbone | Vector type | Bacterial resistance | Addgene reference | Gene insert |
| --- | --- | --- | --- | --- | --- |
| pLJM1-EGFP | pLJM1 | Mammalian expression (3rd generation lentiviral) | Ampicillin 100 µg/mL | #19319 | eGFP |
| pEGFP-N1-TFEB | pEGFP-N1 | Mammalian expression (transient) | Kanamycin 100 µg/mL | #38119 | TFEB-eGFP |

**Table 8:** Non-commercial buffers and solutions.

| Name | Constituents |
| --- | --- |
| 0.4% SRB | 0.4% sulforhodamine B (SRB) in 1% acetic acid |
| Blocking and permeabilization solution | 2% bovine serum albumin (BSA) 0.5% Triton-X in PBS |
| Buffer B | 75% acetonitrile, 20% water, 5% DMSO with + 0.1% ferulic acid (FA) |
| DHB matrix | 20 mg/mL 2,5-dihydroxybenzoic acid (DHB) in 50% acetone with 0.1% trifluoroacetic acid (TFA) |
| Laemmli buffer | 1M Tris, 10% SDS, 10% Glycerol, 5% β-mercaptoethanol, 0.0002% bromophenol blue |
| NBF stop buffer | 101 mM Tris, 100 mM Glycine in DEPC treated water |
| PBS | 137 mM NaCl, 2.7 mM KCl, 8 mM Na<sub>2</sub>HPO<sub>4</sub>, 1.5 mM KH<sub>2</sub>PO<sub>4</sub> |
| Stringent wash | 50% deionized formamide 2X SSC buffer |
| TBS | 50 mM Tris, 150 mM NaCl |
| Tris-glycine running buffer | 2.5 mM Trizma Base, 0.2 M glycine, 0.01% SDS |

Dynamics of rucaparib uptake were captured using eGFP-overexpressing cells (37°C and 5% CO_2_), imaging at 20-minute intervals over 24 hours. Image acquisition was performed as described before. Subsequently, cells were segmented based on their eGFP signal using the Cellpose 2.0 model ‘cyto’, employing flow, cell probability, and stitch thresholds of 0.4, −2, and 0, respectively^50^. The resulting cell masks were then imported into Fiji^42^ using the BioImaging and Optics Platform (BIOP) plugin to quantify rucaparib fluorescence intensity per cell over time. To characterize rucaparib uptake kinetics, a one-phase exponential association or asymptotic growth non-linear model was fitted over the fluorescence intensity values per cell and time point, using GraphPad Prism version 9 (GraphPad Software, San Diego, California USA, Prism).

Rucaparib washout experiments were performed similarly to uptake experiments described above, by pre-staining cells with CellTracker Green and subsequently pretreating with CQ or bafilomycin A1 for 1 hour before rucaparib was added for a further hour. PBS or DMSO were used as vehicle controls for CQ and bafilomycin, respectively. At time 0, drug-containing media was removed and replaced with fresh, drug-free complete RPMI media. Cell tracker green and rucaparib were imaged at 20-minute intervals for 5 hours using the Operetta. The resulting images were analysed in Cellprofiler 4.2.6 to measure average rucaparib per cell, using CellTracker green for cell segmentation. Mean rucaparib fluorescence intensity per cell was then plotted for each of 6 replicate wells per conditions, and a one-phase decay model was fitted in Graphpad prism to calculate half-life per condition.

#### FACS sorting of rucaparib-treated cell populations

Cells were treated with rucaparib at IC_50_ or vehicle only for 1, 2 or 24 hours before washing with PBS, and harvesting using TrypLE express. Cells were resuspended at 5-10 million cells/mL in ice-cold PBS or phenol red-free RPMI, both supplemented with 2% FCS. For experiments requiring re-culture of cells post sorting, 100 U/mL penicillin, 100 µg/mL streptomycin were included in the formulation. Cell suspensions were filtered through a 35 µm cell strainer (Falcon, 352235) and kept on ice prior to sorting on a FACSAria™ Fusion cell sorter (BD Biosciences). Rucaparib was excited by the 355nm laser and emission was collected using a 450/50nm bandpass filter. BD FACS Diva version 9.4 was used for subsequent analyses, and the top and bottom 20% of rucaparib positive cells were sorted and collected into 50% FCS in PBS or phenol red-free RPMI. Samples and collection tubes were maintained at 4°C during sorting and kept on ice prior to downstream processing. For re-culturing, cells were centrifuged at 400 g and resuspended in complete media.

### Molecular biology and biochemistry

#### Lentiviral transduction

HEK293-T cells (grown in DMEM, 100 U/mL penicillin, 100 µg/mL streptomycin, 10% FCS at 37°C, 5% CO_2_), were used for viral particle generation. Cells were transfected using FuGene-6 according to manufacturer’s instructions. Briefly, transfection mix was made by combining serum-free DMEM with FuGene, incubating 5 minutes (room temperature) before adding relevant DNA constructs and further incubating at room temperature (25 minutes). Transfection mix was added dropwise to plates containing HEK293T-cells. After 6 hours, medium was refreshed with complete DMEM and once more the day after transfection. At 48h post transfection, viral particles were harvested by filtering medium using a 0.45 µm filter (Starlab, E4780-1456) prior to adding polybrene (final concentration 4 µg/mL). Virus-containing media was stored at −80°C if not used immediately. For viral transduction, cells of interest were seeded in complete RPMI (approximately 50% confluency), and the following day, were incubated for 6 hours with virus-containing medium (37°C, 5% CO_2_). Post-infection, cells were allowed to recover for 16 hours before puromycin (1 µg/mL).

#### Transient transfection

Cells were seeded in 6-well plates 24h prior to transfection. Cells were transfected with FuGene, using a 1:3 ratio of DNA to transfection reagent, according to manufacturers instructions. Briefly, FuGene-6 (9 µl/well) was added dropwise to serum free medium (138 µl/well) and allowed to incubate for 5 minutes. 3 µg of pEGFP-N1-TFEB plasmid was added to the mix and further incubated for 25 minutes before adding dropwise to the well. Media was replaced after 5-7 hours with fresh complete RPMI, and cells were used for plating after an overnight recovery.

#### Protein extraction and western blotting

Protein was extracted in Laemmli buffer with 100 units/mL benzonase and quantified in a Pierce bicinchoninic acid (BCA) assay (Thermo Fisher, 23227). Typically, 20-40 µg protein were denatured for 5 minutes at 95°C and run per lane in a 4-15% mini-Protean precast protein gels (Biorad, 456-1084) in Tris-glycine running buffer. Protein was transferred to a nitrocellulose membrane (Biorad, 1620112) using the Trans-Blot Turbo Transfer System (Biorad, 15V, 1.5A, 25 minutes) and Biorad protein transfer buffer, stained with ponceau S solution and blocked for 1 hour in 5% milk in TBS-T (0.05% Tween-20 inTris-Buffered Saline). Membranes were then probed with primary antibody (TFEB, D2O7D), diluted 1000 times in 5% BSA in TBS-T with 0.05% sodium azide, at 4°C overnight, followed by 3 consecutive washes in TBS-T and a 1-hour incubation with secondary antibody in 5% milk in TBS-T.

#### Proteomic Liquid chromatography mass spectrometry - sample preparation and analysis

PEO1 cells were treated with rucaparib at IC_50_ concentration or vehicle only, for 1, 2 or 24 hours. Treated cells were FACS-sorted as described before into Ruc_High_ and Ruc_Low_ populations (approximately 1 million cells per replicate), and washed twice with ice-cold PBS before lysis in surfactant cocktail^51^ (2% SDS, 1% SDC, and 2% IGEPAL CA-630), with 100 units/mL benzonase added prior to lysis. Protein was measured by BCA assay and concentration adjusted to 2 g/L. 40 µg of lysate was reduced and alkylated by addition of chloroacetamide and TCEP (tris(2-carboxyethyl)phosphine)) to a final concentration of 20 mM and 10 mM, respectively. Samples were precipitated using 200 µg of beads, washed with 100 µL 80% ethanol three times before digesting overnight with 40 µL of 50 mM ammonium bicarbonate buffer containing 20 ng/µL trypsin (Pierce™ P/N 90059) and 10 ng/µL LysC (WAKO) with shaking (1700 rpm). An Ultimate 3000 RSLC nano liquid chromatography 60 system (Thermo Scientific) was used for chromatographic separation and this was coupled to an Orbitrap HFX mass spectrometer (Thermo Scientific) via an EASY-Spray source. Electro-spray nebulisation was achieved by interfacing to Bruker PepSep emitters (PN: PSFSELJ20, 20 µm). Peptide solutions were injected directly onto the analytical column (self-packed column, CSH C18 1.7µm beads, 300µm × 35cm) at a working flow rate of 5 µL/min for 4 minutes. Peptides were separated using a 66-minute stepped gradient: 0-45% of buffer B (75% acetonitrile, 20% water, 5% DMSO with + 0.1% ferulic acid (FA)) for 66 minutes, followed by column re-conditioning and equilibration. Eluted peptides were analysed by the mass spectrometer in positive polarity, using an initial MS1 at 120,000 resolution followed by sequential MS2 acquisition and ion fragmentation at 30,000 resolution. An m/z range of 409.5 to 1650 was used. Raw Data was processed using Spectronaut (19.0^52^). Initial protein identification (pulsar search) allowed up to three missed cleavages and accounted for common protein modifications. Searches were conducted against the UniProt Homo sapiens 1-gene per protein sequence database (downloaded 22/01/2024, 20,596 entries) and a universal protein contaminants database^53^ (downloaded 22/01/2024, 381 entries). For quantification, MS2-based data was analysed using a Direct DIA approach using the MaxLFQ method^54^, and only proteotypic peptides were considered (q-value < 0.01). After removing entries from the protein contaminants database, further analyses were performed using Perseus (1.6.15.0^55^), where data were log_2_ transformed prior to additional filtering and statistical testing. For two-group comparisons, proteins with at least seven replicate intensities per experimental group were included, and a Student’s t-test was applied with permutation-based FDR correction (q-value threshold of 0.05). Results were visualised as volcano plots, with significance defined by q-value. Perseus Gene Set Enrichment Analysis was performed, where intensities of t-test significant hits were z-score normalised and clustered using Hierarchical Clustering Analysis. Protein IDs within clusters were enriched using the Perseus module for Fisher’s exact test, with multiple-testing correction by Benjamini-Hochberg (q-value threshold of 0.05), assessing the relative enrichment of GO terms per cluster against all background proteins. For Reactome enrichment analysis, UniProt ID mapping service was used to retrieve gene IDs prior to clusterProfiler (4.12.0^56^) analysis, with a q-value threshold of 0.05.

#### Triple Quadrupole Liquid Chromatography-Mass Spectrometry (TQ LC-MS)

For intracellular drug concentration measurements, cells were seeded into 96-well plates at 15000 cells/well and left to adhere overnight. Cells were pre-treated for 1 hour with or without bafilomycin in complete medium before being dosed with PARP inhibitors at their IC_50_ concentrations. After 1 hour of incubation, media were removed and cells were washed once in cold complete medium followed by one wash in cold Dulbecco’s PBS (Gibco, 14190-094). Cells were quenched in cold 80% (v/v) acetonitrile (Honeywell™, Riedel-de Haën™, HPLC gradient grade, 34851-1L) in water (Fisher Scientific, Optima^TM^ LC/MS Grade, 10505904) supplemented with 1.5 ng/mL Amisulpride-d_5_ (CAY30075-1 mg) as an internal standard. Plates were sonicated for 3 minutes before supernatant was transferred to U-bottomed 96-well plates and centrifuged for 1 minute at 1000 rcf. Extracted samples were stored at −80°C. To calculate cell volume, parallel plates of untreated cells were counted using the Nexcelom Cellometer Auto Cell Counter (Nexcelom Bioscience) and cell diameter measurements were recorded.

Samples were thawed on wet ice and diluted 1 in 25 in water (Fisher Scientific, Optima^TM^ LC/MS Grade, 10505904) for analysis using the Waters Xevo® TQ-XS Mass Spectrometer coupled to an ACQUITY Premier System (Waters Ltd, Wilmslow, UK) using a CORTECS T3 Column, 120Å, 2.7 µm, 2.1 mm X 30 mm (Waters, 186008481) at 60°C. A 10 μL volume of sample was injected. Mobile phase A1 contained water (Fisher Scientific, Optima^TM^ LC/MS Grade, 10505904) with 0.1% (v/v) formic acid and mobile phase B1 contained acetonitrile with 0.1% (v/v) formic acid (LC-MS Ultra, Honeywell™ Riedel-de-Haën™ CHROMASOLV™, 15611410). A linear gradient from 1% to 100% mobile phase B1 was applied over 3 minutes at a flow rate of 1.2 mL/min. The sample eluate was injected into an XEVO® TQ-XS mass spectrometer using electrospray ionisation in positive ion mode. MS conditions: source temperature 150 °C; capillary voltage 0.8 kV; desolvation temperature 600°C; desolvation gas flow 1000 L/Hr; cone gas flow 150 L/Hr. System settings were controlled using MassLynx 4.2 software (Waters Ltd, Wilmslow, UK) and multiple reaction monitoring methods were optimised using IntelliStart and validated manually by direct infusion. Cone voltages (V) and collision energies (eV) were optimised for quantifier and qualifier mass transitions as follows: Amisulpride-d_5_ quantifier 375.468 > 242.251 *m/z* (38 V, 28 eV), qualifier 375.468 > 196.237 *m/z* (38 V, 40 eV), retention time 0.86 min; rucaparib quantifier 323.984 > 293.05 *m/z* (34 V, 10 eV), qualifier 323.984 > 235.962 *m/z* (34 V, 32 eV), retention time 0.935 min; Niraparib quantifier 321.13 > 205.11 *m/z* (8 V, 42 eV), qualifier 321.13 > 304.14 *m/z* (8 V, 18 eV), retention time 1.0 min; Olaparib quantifier 435.18 > 281.3 *m/z* (10 V, 30 eV), qualifier 435.18 > 367.3 *m/z* (10 V, 20 eV), retention time 1.165 min. MassLynx 4.2 software automatically calculated the dwell time for each transition with a minimum of 15 analytical points per peak. Samples were run alongside matrix-matched calibration curves generated using untreated cell extracts processed in parallel to the treated samples.

Data were processed using Skyline software^57^. Peak areas were normalised to the internal standard and plotted against the calibration curve before values were normalised to cell density measurements so intracellular concentration was calculated. Values were normalised to PARP inhibitor only controls and data were plotted using GraphPad Prism version 9 (GraphPad Software, San Diego, California USA, Prism).

## Acknowledgements

The authors would like to thank Hiromi Kudo and Robert Goldin for help with immunohistochemistry, Anke Nijhuis for providing cell lines, Nazma Malik for sharing the TFEB-GFP transient expression construct, Ka Lok Choi for assistance with genomics approaches, James Elliot and Joana de Teixeira Carrelha for assistance with flow cytometry, Oliver Gonzalez Carvajal for assistance with cell collection and Aleksandra Gruevska for advice and assistance in mass spectrometry imaging. We would also like to thank the patients who gave their consent for their material to be used in this study. Tumour tissues samples were provided by the Imperial College Healthcare NHS Trust Tissue Bank. Other investigators may have received samples from these same tissues. The research was supported by the National Institute for Health Research (NIHR) Biomedical Research Centre (BRC) based at Imperial College Healthcare NHS Trust and Imperial College London. The views expressed are those of the author(s) and not necessarily those of the NHS, the NIHR or the Department of Health. C.R.M. was supported by funding from the Integrative Toxicology Training Partnership, administered via the MRC Toxicology Unit, awarded to L.F. and Z.H. Z.H. acknowledges funding from the MRC (MR/W019132/1). This work was also supported by intramural funding from the Medical Research Council (MC-A564-5QC70) and a CRUK Career Establishment Award (RCCCEA-Nov21\100001) awarded to L.F.

## Author Contributions

Conceptualisation, C.R.M. and L.F.; Methodology, C.R.M, L.F., P.C., Z.H. and A.R.B; Formal Analysis, C.R.M., G.Z., R.R., G.R.Y. A.M., D.M. and L.F.; Investigation, C.R.M., G.Z., P.O.P., E.D. and N.L.; Resources, C.W., I.A., B.P., B.R.P., V.W., Z.T., N.M., P.S., L.G., B.L., I.M.,C.F. and A.R.B. ; Writing (original draft), L.F.; Writing (review and editing), L.F., C.R.M., Z.H., P.C., C.F., B.L. and A.R.B; Supervision, L.F. and Z.H.; Funding Acquisition, L.F. and Z.H.

## Extended Data Figures

***Extended Data Figure 1 (associated with main* *Figure 1*)**

A) Schematic representation of mass spectrometry imaging calibration line generation process in Atmospheric-Pressure Matrix-Assisted Laser-Desorption ionisation (AP-MALDI) and Desorption Electrospray Ionisation (DESI) platforms. Calibrants were spotted on top of un-dosed HGSOC tissue sections.

B) Adducts of niraparib, olaparib and rucaparib detected in AP-MALDI platform, in positive ion mode with 2,5-dihydroxybenzoic acid (DHB) matrix. Total Ion Chromatogram (TIC)-normalisation and a tolerance window of 2.5 parts per million (ppm) was applied.

C) AP-MALDI (left) and DESI (right) calibration lines generated in positive or negative (DESI, rucaparib only) ion modes.

D) PDE dosing timepoint optimisation schematic. Tissue explants were dosed with olaparib at 20 µM for up to 48 hours to determine time required for the compound to homogeneously diffuse through the top 150 µm of tissue. This depth was selected to match the number of 10 µm sections typically used in downstream analyses, ensuring that drug exposure reached steady state across all sections.

E) For each sample, the 6^th^ and 12^th^ section (72 and 144 µm deep respectively) were thaw-mounted in a glass slide for spectral data acquisition using DESI in positive ion mode. Comparison between tissue architecture as seen through haematoxylin and eosin (H&E) staining (scale bar = 750 µm) and olaparib signal throughout time is shown ([M+Na]^+^ adduct, experimental m/z 457.1670, mass accuracy = −3.96 ppm).

F) Quantification of olaparib in 6^th^ and 12^th^ sections throughout time, calculated as the average TIC-normalised ion intensity value per tissue section area, thresholded through haematoxylin and eosin staining optical density.

***Extended Data Figure 2 (associated with main* *Figure 1*)**

2-part figure depicting serial sections from all treated PDEs used to perform drug MSI signal analysis, haematoxylin and eosin (H&E) staining, immunoistochemical (IHC) analyses, and where relevant, GeoMx spatial transcriptomics, where chosen ROIs are shown. Key indicating colour codes for patient, tumour site and drug in part 2 of figure. For IHC analyses (rows 1, 2, 9 and 10) pseudo-colour overlays highlight positive cells compared to negative controls (no primary antibody).

***Extended Data Figure 3 (associated with main* *Figure 2*)**

A) Regions of interest (ROI) included in GeoMx spatial transcriptomics study. On average, 5 ‘high-‘ and 5 ‘low-drug’ ROIs were chosen across PDE samples from three patients, including technical replicates when available, resulting in approximately 45 ROIs per compound and drug content. Per compound, the total number of ROIs obtained is shown on the left, with a breakdown by patient displayed on the right, before and after data quality control (QC) and filtering. QC involved applying a threshold of 100,000 raw reads and 50% sequencing saturation, discarding any ROI that fell below these criteria. ROIs with less than 5% of targets above the limit of quantification were filtered out.

B) Supervised hierarchical clustering of GeoMx spatial transcriptomics dataset. Samples were grouped by Patient ID before hierarchical clustering analysis performed using Euclidean distance. All 11224 targets are shown.

C) Principal component analysis of niraparib and rucaparib datasets. ROIs are coloured based on drug content, with outlines representing patient clusters.

***Extended Data Figure 4 (associated with main* *Figure 2*)**

A) Volcano plot depicting genes identified in niraparib linear mixed model regression analysis. Genes significantly associated with increasing drug illustrated in grey points (p < 0.05), genes illustrated with red points (and text) are also significant in the rucaparib analysis.

B) Volcano plot depicting genes identified in rucaparib linear mixed model regression analysis. Genes significantly associates with increasing drug illustrated in grey points (p < 0.05), genes illustrated with red points (and text) are also significant in the niraparib analysis.

C) Gene set enrichment analysis (GO biological processes) for genes significantly associated with increasing levels of niraparib (left) and rucaparib (right) within PDE samples in linear mixed model regression analysis.

D) Scatter plots depicting expression of lysosomal genes relative to niraparib concentration from PDE experiments, patients represented in different colours and Pearson correlation coefficients indicated.

***Extended Data Figure 5 (associated with main* *Figure 3*)**

A) Dose response curves to all three PARP inhibitors (niraparib, olaparib and rucaparib) for 6 OvCa cell lines, measured by SRB assays. Data represents n = 3 pooled biological replicates (each with n = 3 technical replicates), dosed with 10 concentrations ranging from 0.95 to 100uM for 72 h.

B) IC_50_ values of OvCa cell lines measured by SRB assays. Estimated values by fitting appropriate non-linear regression model in Graphpad Prism: [Inhibitor] vs. response – Variable slope (four parameters) for all lines except Kuramochi, which was fitted with the [Inhibitor] vs. normalized response model

C) Heterogeneity in response to PARP inhibitors measured by γH2AX level in OvCa cell lines. Each cell line was dosed at their corresponding IC_50_ values for each PARP inhibitor for 24h. P-values shown for Welch two sample t-test comparing mean γH2AX per technical replicate (n = 3) in PARP inhibitor-treated cells compared with vehicle control. Data representative of n = 3 biological replicates.

***Extended Data Figure 6 (associated with main* *Figure 3*)**

A) eGFP-PEO1 treated with vehicle-only or rucaparib at IC_50_ concentration for 24 hours, 100 μm scale bar.

B) Live PEO1 single-cell rucaparib accumulation kinetic parameters, estimated with a one-phase association non-linear model, fitted over mean rucaparib fluorescence intensity per cell over time. Pearson’s correlation coefficient between rate constant K and maximum rucaparib per cell was calculated over 3 colour-coded independent biological replicates. Black line marks the average values.

C) Ruc_High_ and Ruc_Low_ PEO1s (top and bottom 20% of the rucaparib-treated population respectively) were FACS-sorted with total rucaparib and vehicle-only controls after a 2-hour treatment. Cells were re-cultured in drug-free medium for 7 and 14 days and then re-dosed to analyse rucaparib signal with respect to its original distribution (left). Relative drug levels for each population are shown on the right. ‘First’ denotes initial rucaparib signal. ‘Second’ refers to re-dosed levels after re-culture. Data representative of n = 3 technical replicates.

***Extended Data Figure 7 (associated with main* *Figure 4*)**

A) Median drug signal intensity per FACS gate in PEO1 cells treated and sorted based on rucaparib fluorescence intensity. Rucaparib signal was normalised to the signal intensity of the Rucaparib-treated, ungated population (Ruc total) per time point and biological replicate (n = 4). Two technical teplicates per biological replicate are depicted.

B) PEO1 cell volume per FACS gate. Cell volume was estimated through median FSC-A per gate, which is proportional to cell diameter. Volumes were normalised, per biological replicate, to that of 1 hour vehicle-treated cells.

C) Rucaparib concentration per FACS gate of PEO1 cells. Rucaparib signal intensity was normalised to cell volume, estimated through FSC-A as a proxy of cell diameter. For A-C, two-way ANOVA with the Geisser-Greenhouse correction and Tukey’s or Sidak multiple comparison test were applied to evaluate statistical significance.

D) Example of colocalisation of rucaparib with the lysosome at 60 minutes post dosing in OVCAR4 cells. Scale bar = 20 µm.

E) FACS-based correlation between lysosomal and rucaparib contents per PEO1 cell, after 1.5h dosing of both compounds (left). Cells were gated as before (see Fig. 4a), and median LysoTracker signal per gate was plotted relative to each biological replicate’s total signal (right).

F) Western blot to demonstrate TFEB-GFP overexpression in PEO1 cells.

G) Left; quantification of rucaparib signal per cell in untransfected PEO1 cells, or cells transfected with TFEB-GFP to induce lysosomal biogenesis. Data representative of n = 3 biological replicates. Right; median rucaparib levels per biological replicate (n = 3). Welch two sample t-test was applied to assess significance.

H) Left; quantification of rucaparib signal per PEO1 cell following CQ treatment. Data representative of n = 3 biological replicates. Right; median rucaparib levels per biological replicate (n = 3). Welch two sample t-test was applied to assess significance.

I) Left; quantification of rucaparib signal per PEO1 cell following BafA1 treatment. Data representative of n = 3 biological replicates. Right; median rucaparib levels per biological replicate (n = 3). Welch two sample t-test was applied to assess significance.

J) Left; quantification of rucaparib signal per OVCAR4 cell following BafA1 treatment. Data representative of n = 3 biological replicates. Right; median rucaparib levels per biological replicate (n = 3). Welch two sample t-test was applied to assess significance.

***Extended Data Figure 8 (associated with main* *Figure 5*)**

A) Schematic representation of palbociclib mechanism of action.

B) PEO1 cells were treated with palbociclib or vehicle-only for 24 hours and cultured in drug-free medium for an additional 24 hours before fixation and DNA staining with hoechst. Cell cycle gating was performed based on hoechst sum fluorescence intensity to estimate percentage of cells in each cell-cycle phase.

C) The G2/M gate of the palbociclib-treated OVCAR4 cell population was subdivided into quartiles based on nuclear rucaparib signal to compare γH2AX levels between groups. Data representative of n = 3 biological replicates.

D) In OVCAR4 cells, median γH2AX is linearly related to median intracellular rucaparib within the quartile. Data representative of n = 3 pooled biological replicates.

***Extended Data Figure 9 (associated with main* *Figure 6*)**

A) Left; quantification of nuclear levels of rucaparib (per PEO1 cell), with or without treatment with bafilomycin. Right; correlation between nuclear and cytoplasmic drug levels with or without pre-treatment with bafilomycin (p < 0.05). Data representative of n = 3 biological replicates.

B) Median nuclear rucaparib per PEO1 biological replicate treated with bafilomycin (n = 3). Welch two sample t-test was applied to assess significance.

C) Left; quantification of nuclear levels of rucaparib (per PEO1 cell), with or without treatment with EN6. Right; correlation between nuclear and cytoplasmic drug levels with or without pre-treatment with bafilomycin (p < 0.05). Data representative of n = 3 biological replicates.

D) Median nuclear rucaparib per PEO1 biological replicate treated with EN6 (n = 3). Welch two sample t-test was applied to assess significance

E) Representative images of rucaparib in PEO1 cells after 1 hour with or without pre-treatment with chloroquine (CQ). Dotted white lines outline the nucleus. Scale bar represents 50 µM.

F) Left; quantification of nuclear levels of rucaparib (per PEO1 cell), with or without treatment with chloroquine. Right; correlation between nuclear and cytoplasmic drug levels with or without pre-treatment with chloroquine (p < 0.05). Data representative of n = 3 biological replicates.

G) Median nuclear rucaparib per PEO1 biological replicate treated with chloroquine (n = 3). Welch two sample t-test was applied to assess significance.

H) Representative images of rucaparib in OVCAR4 cells after 1 hour with or without pre-treatment with bafilomycin (BafA1). Dotted white lines outline the nucleus. Scale bar represents 50 µM.

I) Left; quantification of nuclear levels of rucaparib (per OVCAR4 cell), with or without treatment with bafilomycin. Right; correlation between nuclear and cytoplasmic drug levels with or without pre-treatment with bafilomycin (p < 0.05). Data representative of n = 3 biological replicates.

J) Median nuclear rucaparib per OVCAR4 biological replicate treated with bafilomycin (n = 3). Welch two sample t-test was applied to assess significance.

K) Rucaparib fluorescence loss over time in PEO1 cells after washout, with or without chloroquine. Data shows mean rucaparib fluorescence intensity per cell ± standard deviation calculated across 6 replicate wells and is representative of 2 independent biological replicates.

L) Intracellular concentration of olaparib, niraparib and rucaparib in OVCAR4 cells with or without treatment with bafilomycin to alkalinise lysosomes. Data represents combined data from 2 biological replicates, each within 4 technical replicates, and plotted relative to PARP inhibitor only control, within biological replicate. Statistical significance assessed by 2-way ANOVA with Dunnet’s correction for multiple comparisons.

## Notes

**Declaration of Interests:** The authors declare no conflicting interests.

### Competing Interest Statement

IM has undertaken advisory boards for AstraZeneca, Clovis Oncology/pharma& and GSK. Imperial College has received institutional grant support from AstraZeneca

