## Supplementary Figures for "Multimodal imaging reveals a lysosomal drug reservoir that drives heterogeneous distribution of PARP inhibitors"

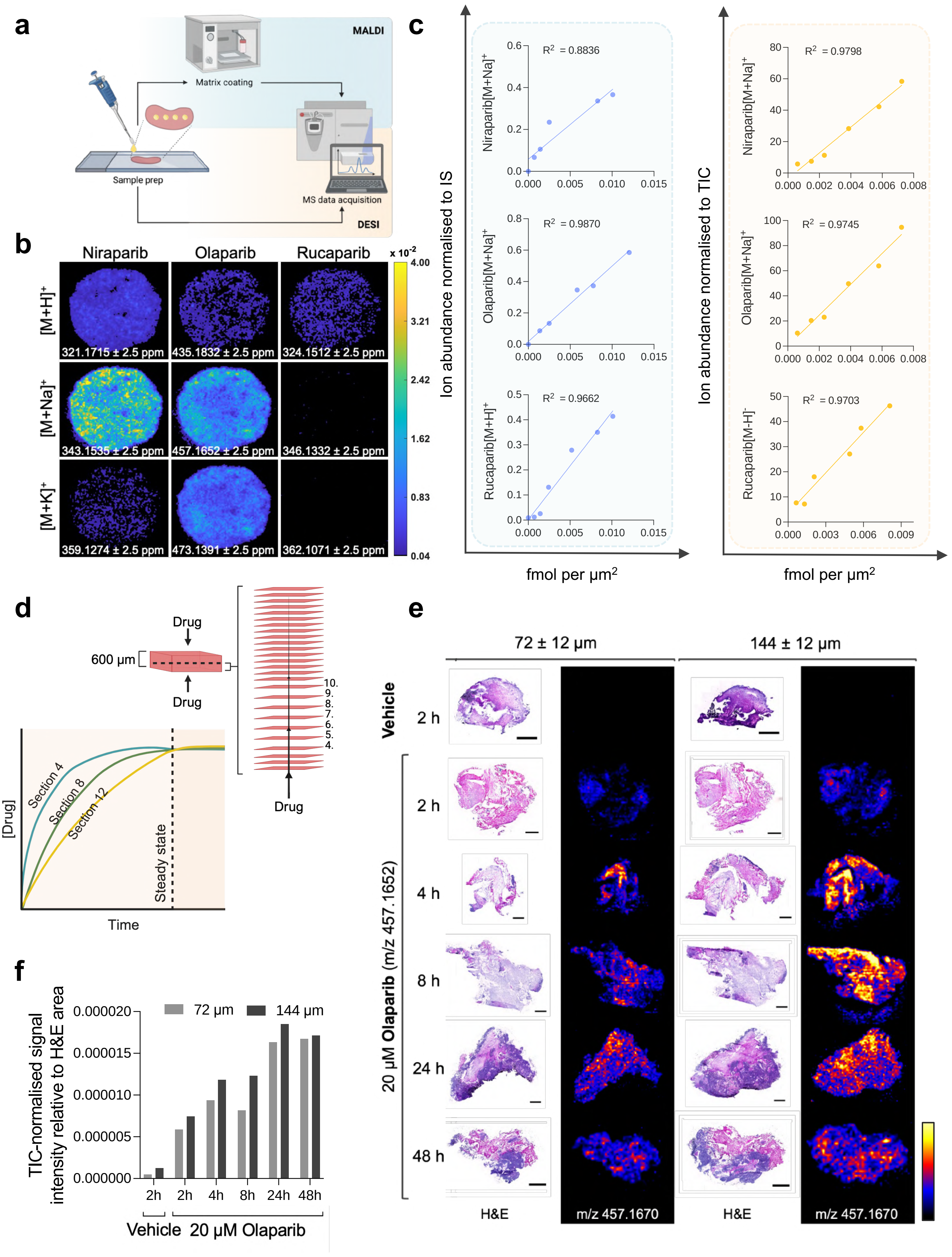


### Extended Data Fig 1

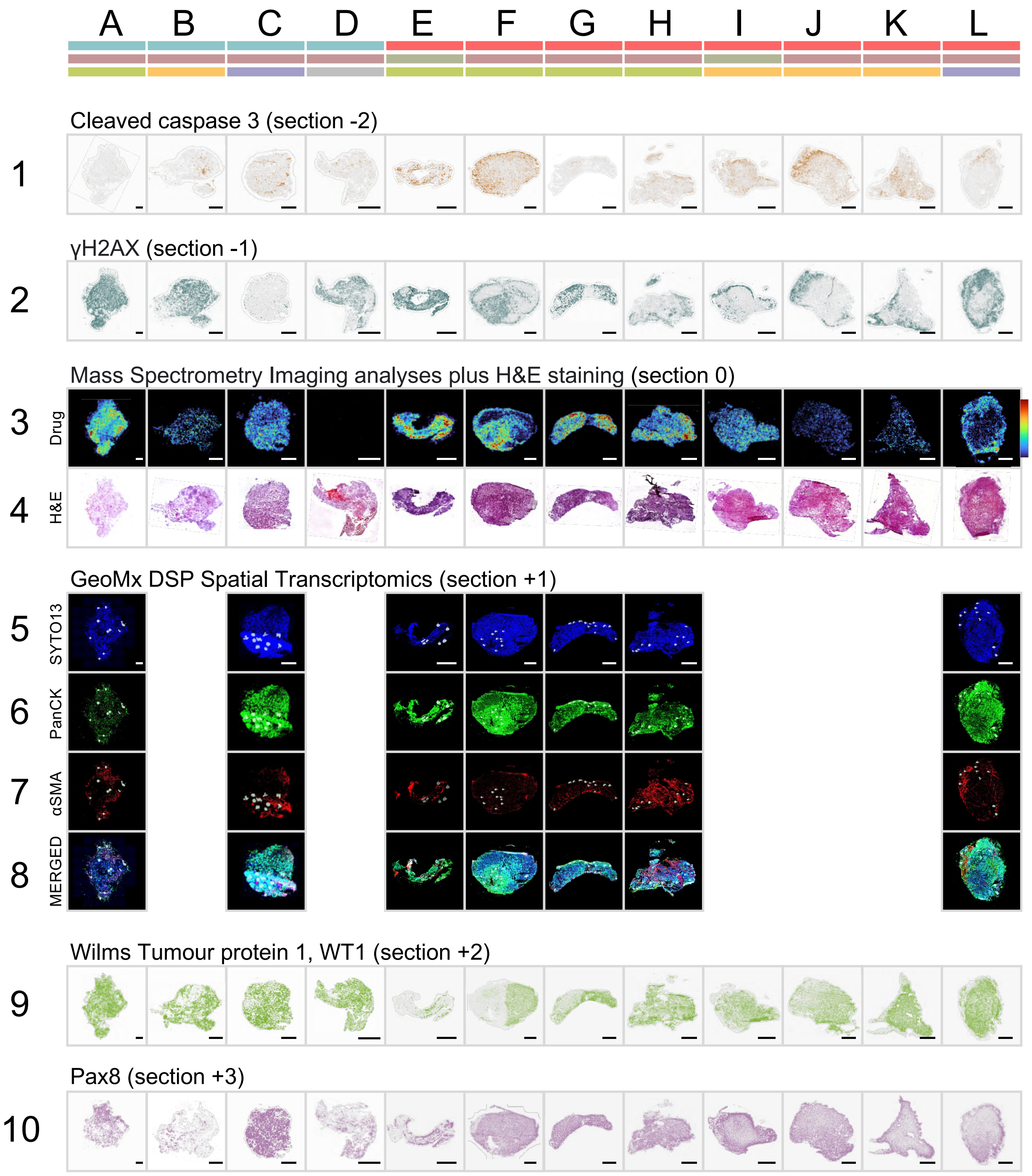


Extended Data Fig 2

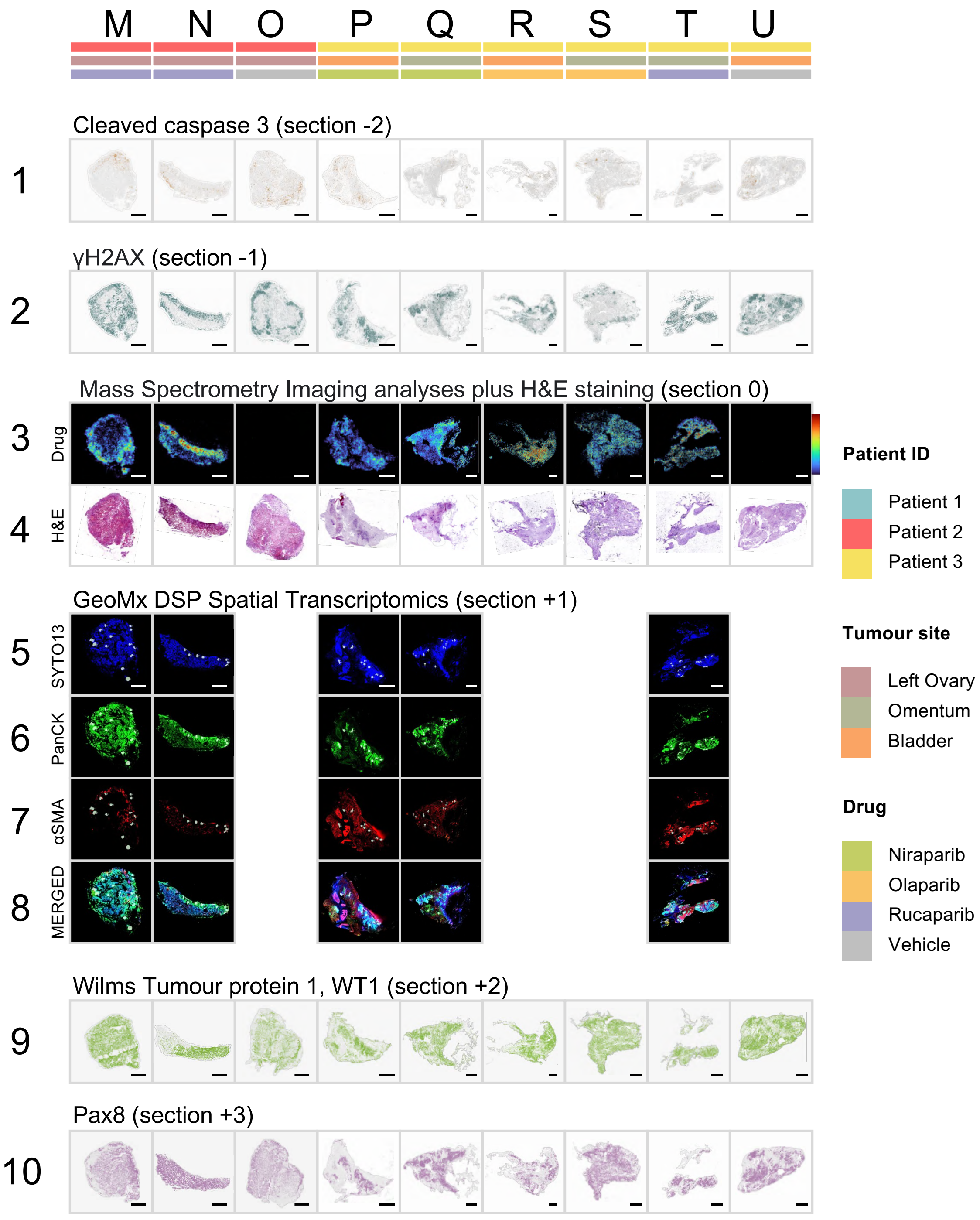


Extended Data Fig 2 continued

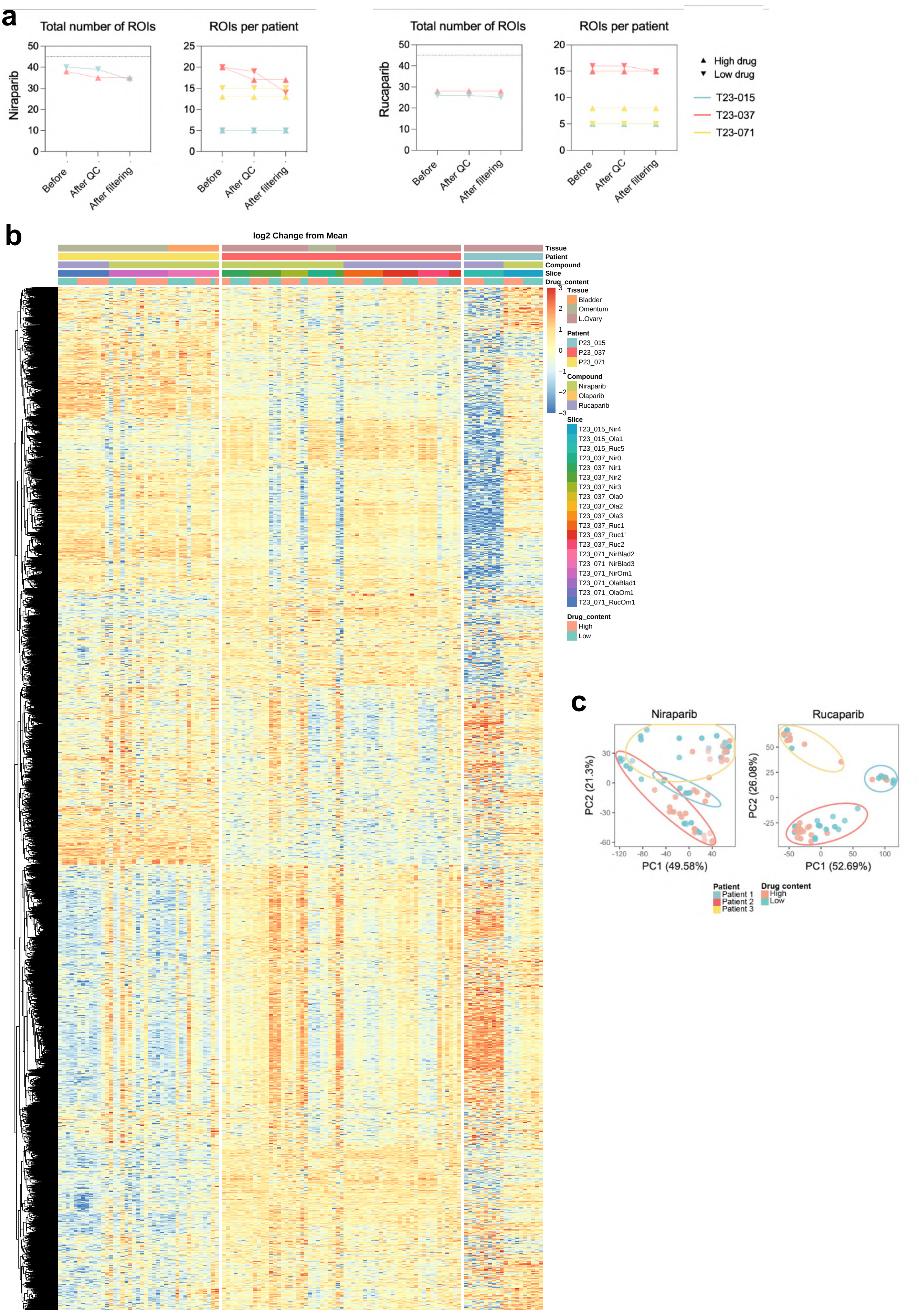


Extended Data Fig 3

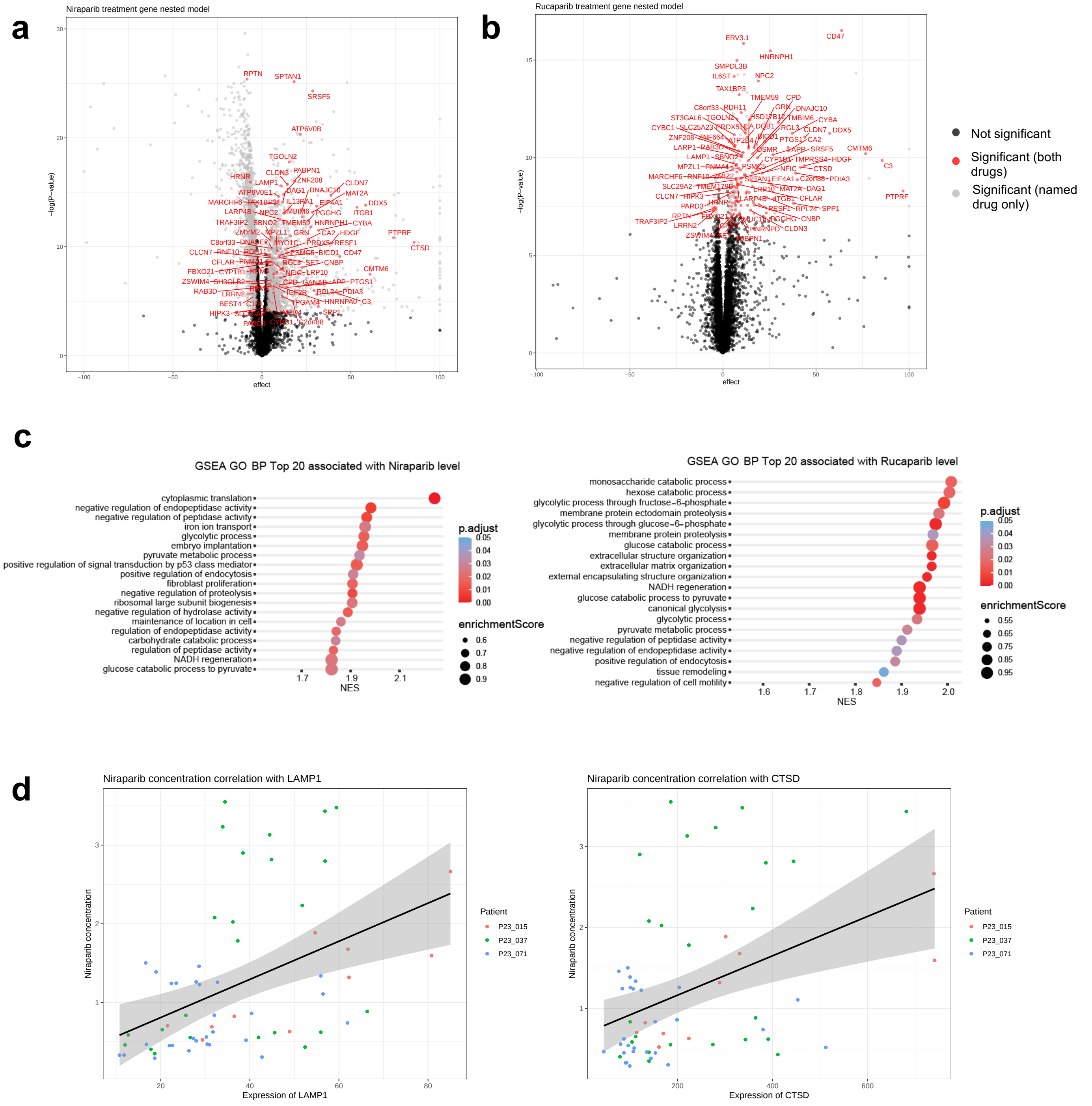


| Gene | Pearson rho | *P* Value |
|---|---|---|
| LAMP1 | 0.442 | 3.2e-4 |
| CTSD | 0.421 | 6.5e-4 |

### Extended Data Fig 4

a

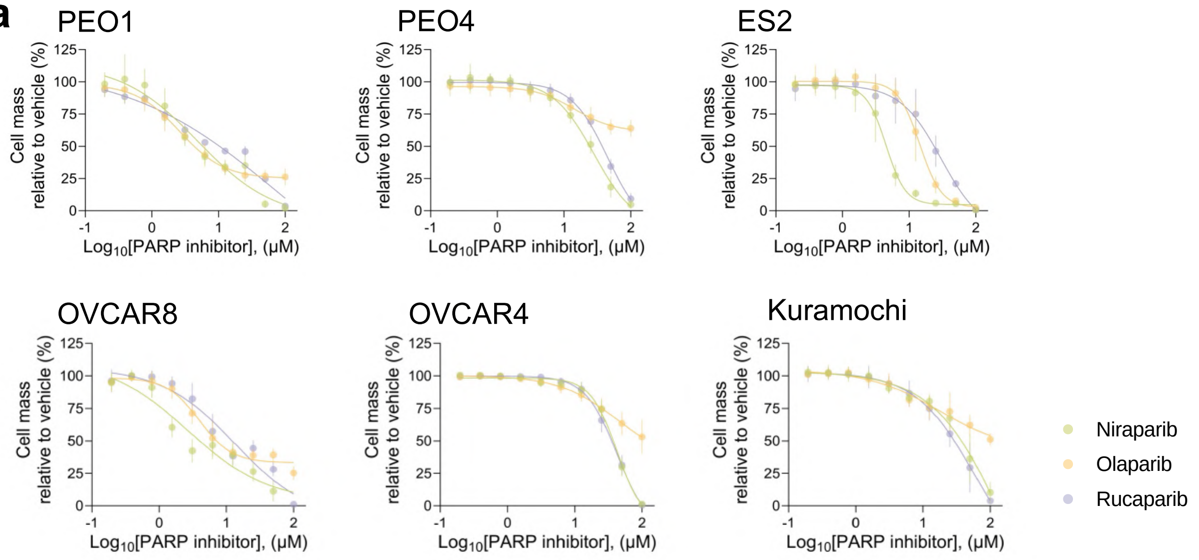


b

| IC50 (μM) | Niraparib | Olaparib | Rucaparib |
|---|---|---|---|
| PEO1 | 6.5 | 10 | 8.5 |
| PEO4 | 38 | 15 | 40 |
| ES2 | 4.5 | 14.5 | 30 |
| OVCAR8 | 2.5 | 4 | 12.5 |
| OVCAR4 | 42 | 38.5 | 42 |
| Kuramochi | 34 | 17.5 | 26 |

c

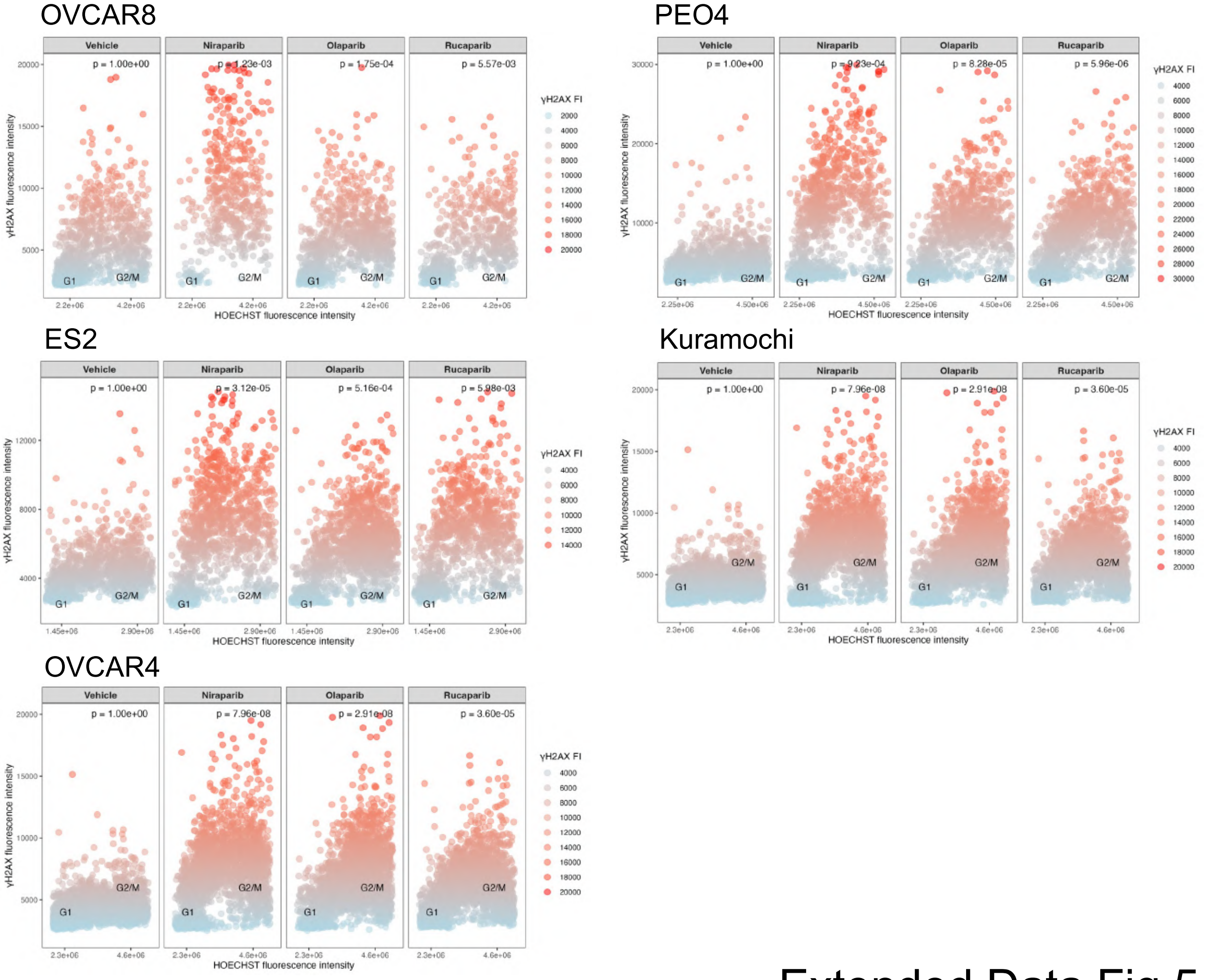


Extended Data Fig 5

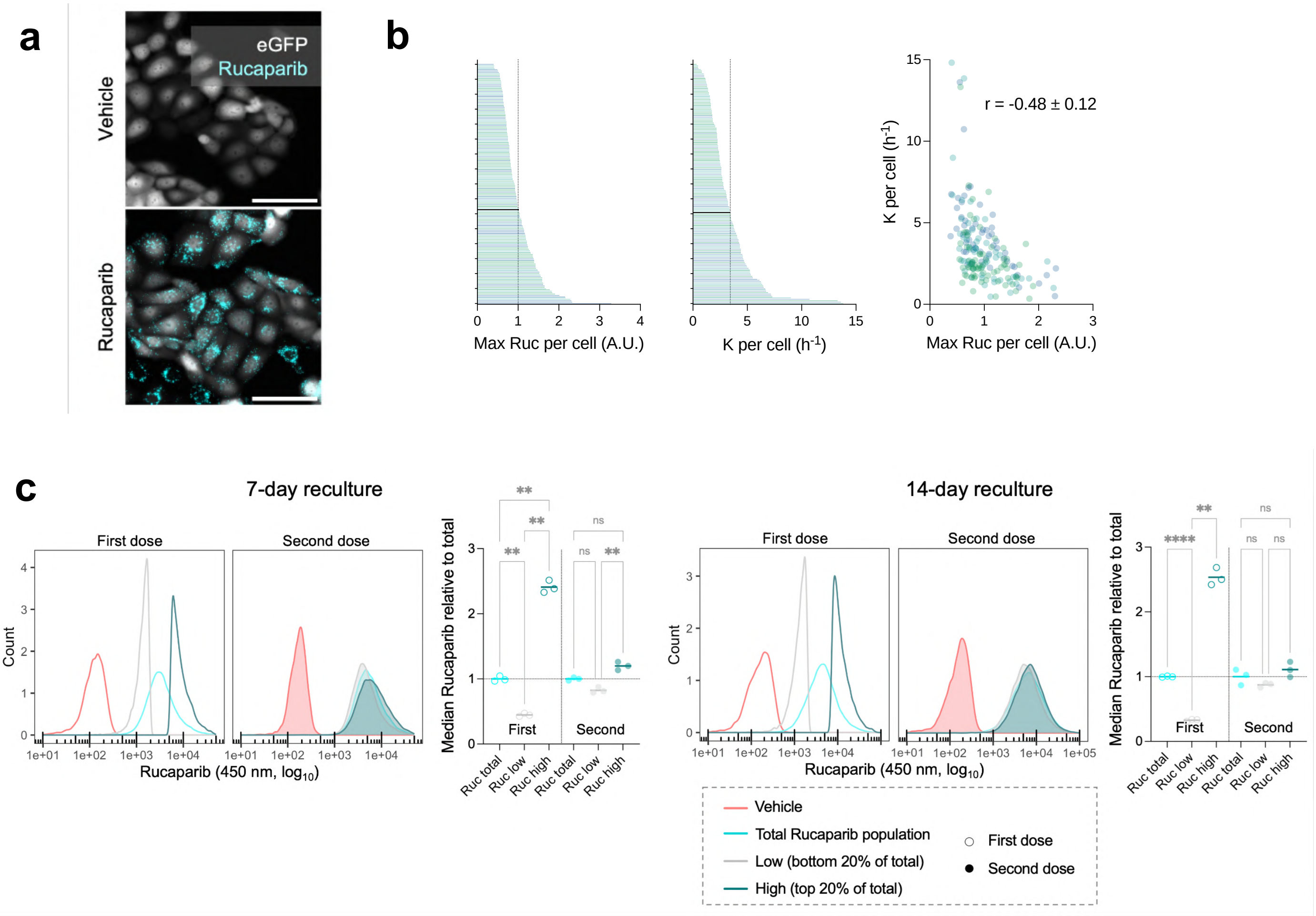


Extended Data Fig 6

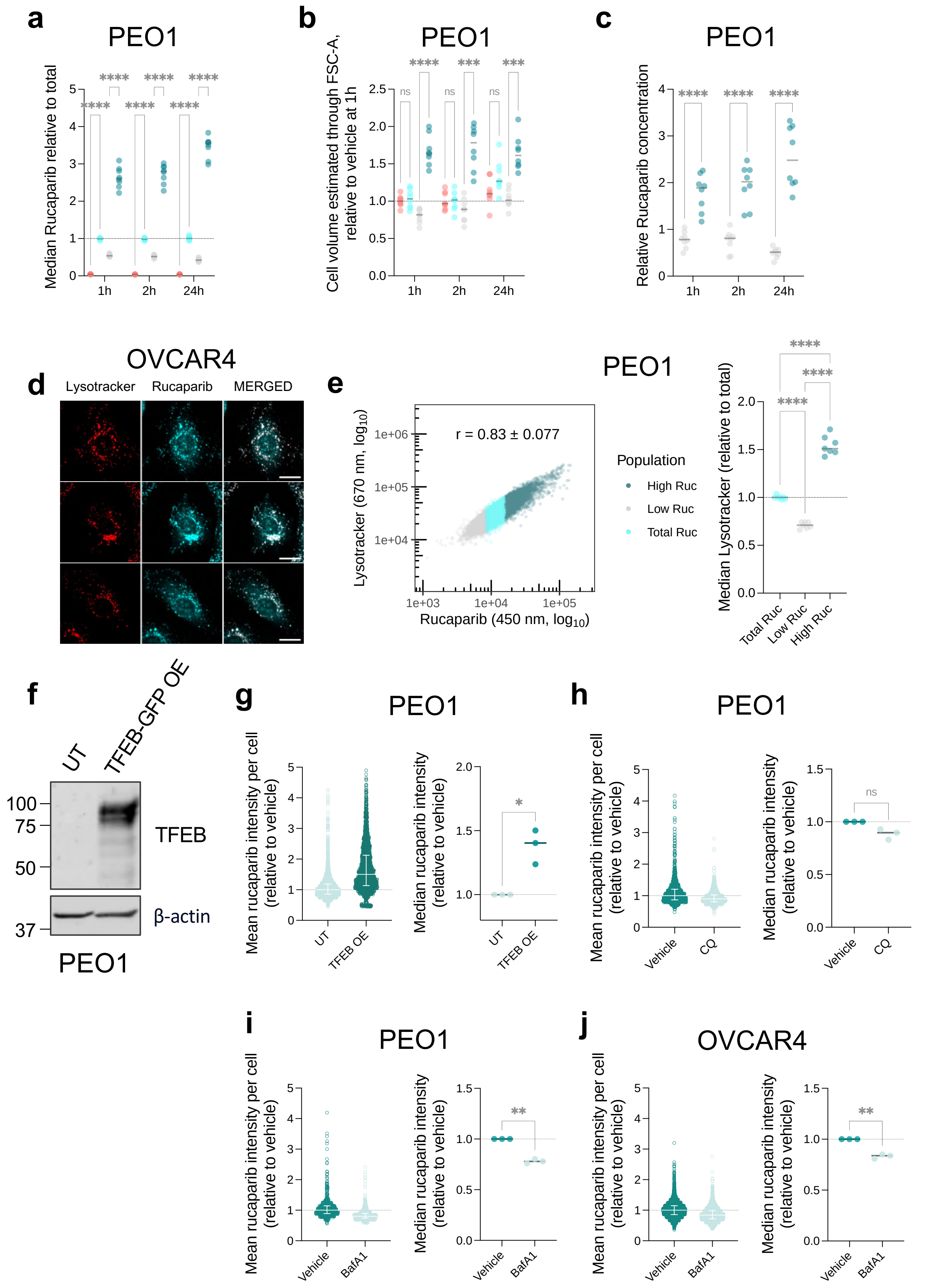


### Extended Data Fig 7

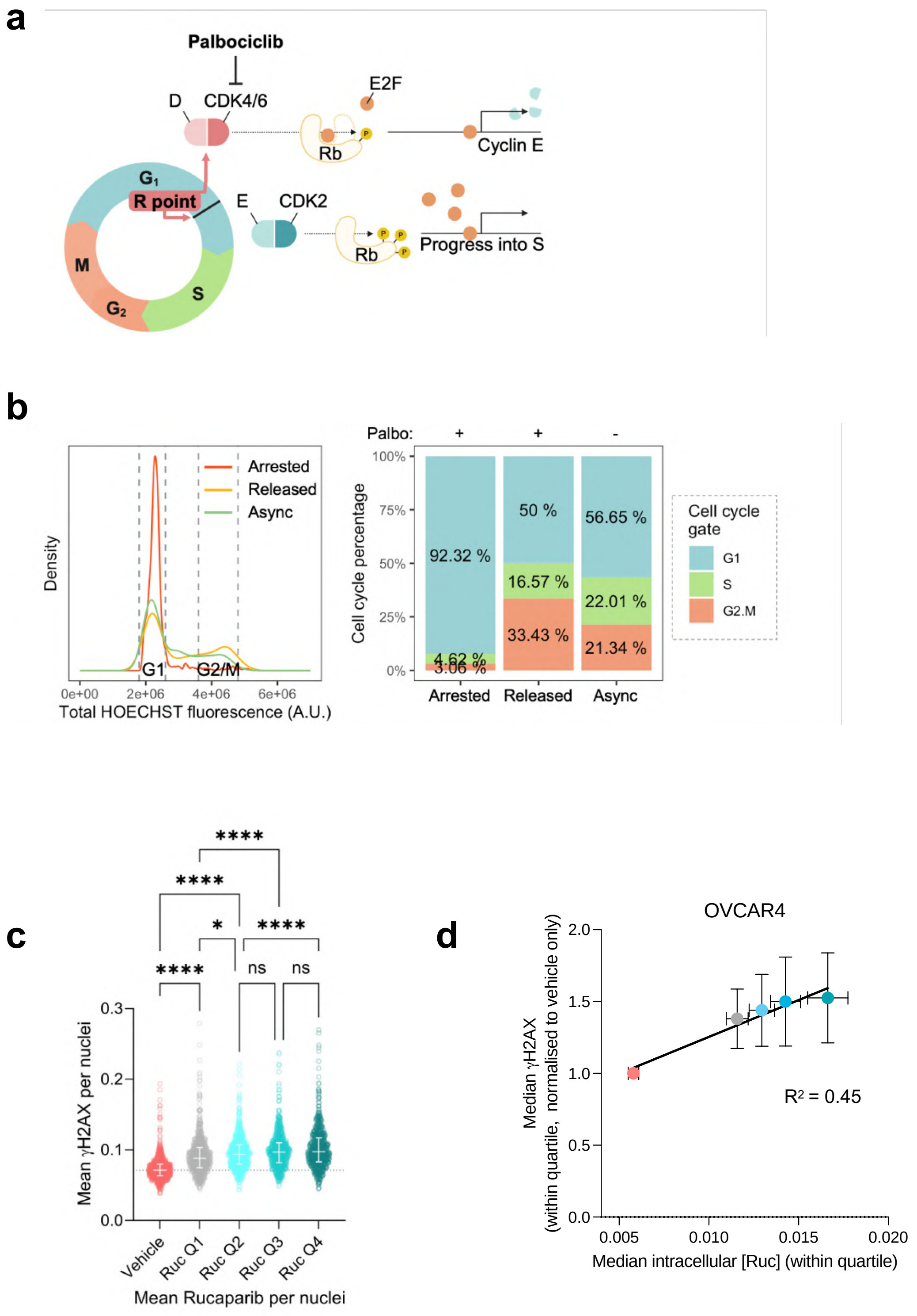


Extended Data Fig 8

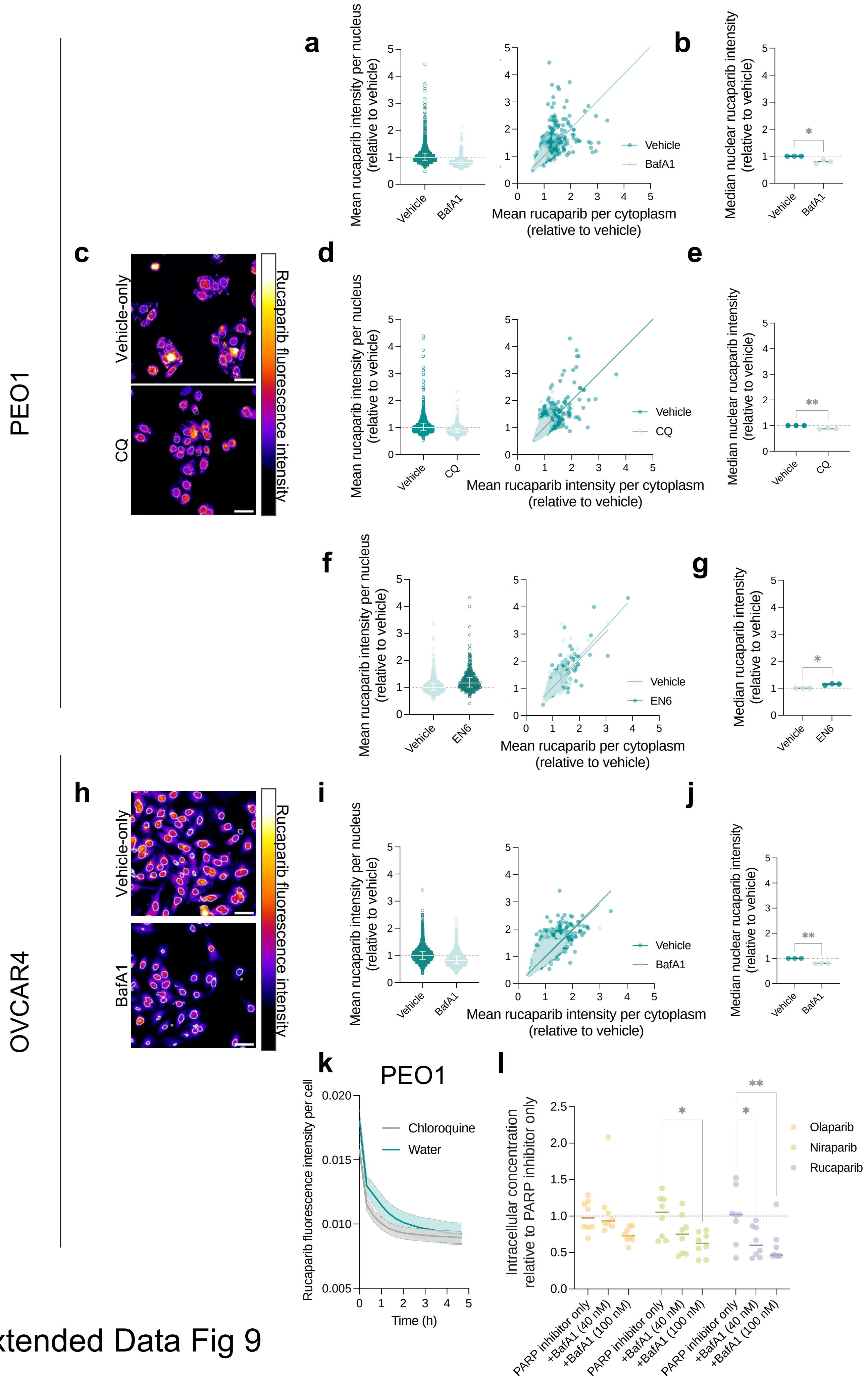


Extended Data Fig 9
